# Transcriptome analysis of *Diplolepis rosae*: revealing overexpression of genes potentially associated with insect immune response and gall formation at early larval stages

**DOI:** 10.1101/2023.11.06.564334

**Authors:** Ksenia Mozhaitseva¹, Zoé Tourrain¹, Antoine Branca¹

## Abstract

Insect parasites can provoke drastic changes in host plant physiology by affecting cell differentiation pathways and various metabolic processes. An intriguing example of such interaction is a gall, a novel outgrowing plant organ, induced by another organism for its own benefit. Cynipidae is a family of gall-inducing hymenopterans that induce galls with a complex anatomical structure. Gall formation involves three stages: initiation, growth, and maturation. Until today, the mechanism of gall initiation remains unknown. In this study, we aimed to reveal candidate genes involved in gall induction in *Diplolepis rosae*, a gall wasp inducing bedeguars in wild roses. We performed differential expression analysis of the gall wasp larva transcriptome. We observed the overexpression of genes encoding plant cell wall degrading enzymes. These enzymes may contribute to the formation of a chamber for a developing larva by lysing plant tissues. We also demonstrated the overexpression of genes encoding podocan, vasorin-like protein, toll-like receptor 7, tetraspanin, lipase, peroxidase, phospholipase A2, and venom acid phosphatase. These genes may be involved in insect development and the immune response against parasitoids, host plant microbiome, and host plant defense systems. Additionally, we performed a test for selection to detect *D. rosae* genes under positive selection. However, we detected only one gene encoding a transposable element. This can be explained by the mostly asexual reproductive mode of this species. Thus, our study contributes to understanding the processes occurring in cynipid wasps during gall formation and creates opportunities for further investigations of other candidate genes.

## Introduction

In host-plant - parasite interactions, parasites can manipulate their host plants through cell reprogramming, metabolic alterations, and immune system suppression. A fascinating example of such an interaction is the formation of galls, abnormal plant outgrowths induced by foreign organisms for their own benefit (Gätjens-Boniche, 2019). Gall-inducing organisms use galls as a nutritional resource and protection against unfavorable biotic and abiotic conditions (Stone & Schönrogge, 2003).

Gall formation can be caused by both micro- and macroorganisms from different taxa (Gätjens-Boniche, 2019). However, the fine molecular mechanism underlying gall formation is well understood only for a handful of microorganisms like *Sinorhizobium* spp. inducing root nodules in Fabaceae, *Agrobacterium tumefaciens* provoking crown galls, and *Ustilago maydis*, the smut fungus of maize (Le Fevre et al., 2015; Hearn et al., 2019). In these microorganisms, the mechanism of gall formation includes (1) detection of released host plant molecules by the infectious agent, (2) release of gall inducer effector molecules interacting with specific plant receptors, and (3) further induction of plant growth response resulting in the formation of gall tissue (Le Fevre et al., 2015).

Unlike microorganisms, gall formation by animals is more complex and less understood, except for a few economically important invertebrates such as root-knot and cyst nematodes (Mejias et al., 2019), several aphids (Nabity et al., 2013; Korgaonkar et al., 2021), and the Hessian fly, *Mayetiola destructor* (Stuart et al., 2012). Invertebrate gallers are shown to induce changes in sugar and nitrogen metabolism, disruption of the defense system, and cellular modifications in host plants (Giron et al., 2016). However, little is known about triggers encoded in the genomes of gall-inducing invertebrates, such as insects, that could be responsible for gall initiation. Unlike gall-inducing microorganisms, whose activity leads to the formation of unstructured tumor tissue in plants (Gätjens-Boniche, 2019), and herbivore animals that simply damage plant tissues, insect gall inducers employ mechanisms that cause significant alterations in host plant metabolism and reprogram cell differentiation, ultimately resulting in the development of a structured new organ. It is fascinating how such a complex phenotype as the insect gall can be initiated by molecular triggers originating from the insect beyond its body. The genetic basis, i.e. the genes encoding the molecular triggers that provoke gall formation in insect gallers, remains a relevant question.

Today, multiple hypotheses, that do not mutually exclude one another, propose potential molecular triggers for gall formation. Firstly, gall initiation is believed to occur at the moment of oviposition, suggesting that components detected in female venom glands and egg secretions could serve as potential triggers (Cambier et al., 2019; Gobbo et al., 2020). Secondly, salivary gland secretions of developing larvae may contribute to gall formation (Zhao et al., 2015; Hearn et al., 2019; Korgaonkar et al., 2021). Thirdly, as gallers manipulate plant cell differentiation and metabolic pathways, they are suggested to produce various molecules that mimic phytohormones (Yamaguchi et al., 2012). Lastly, we can also consider the role of symbiotic or pathogenic microorganisms or microbial genes acquired through horizontal gene transfer, which may also play a part in gall initiation (Bartlett & Connor, 2014; Hearn et al., 2019).

The most studied genes potentially involved in galling are those of *M. destructor* (Diptera: Cecidomyiidae), one of the most destructive crop pests. More than 7% of the Hessian fly genome has been estimated to encode putative effector proteins. This group of proteins includes secreted salivary gland protein (SSGP)-71. This protein contains leucine-rich repeats (LRR) that mediate protein-protein interactions (Ho et al., 2006). Moreover, in plants, proteins containing LRRs play a role in plant development and immunity (Zhao et al., 2015). Interestingly, the whole structure of (SSGP)-71 resembles ubiquitin E3 ligases in plants and E3-ligase-mimicking effectors in plant pathogenic bacteria. SSGP-71 protein has been shown to interact with wheat Skp signal protein *in vivo*. Mutations in the gene encoding SSGP-71 have been shown to avoid the effector-triggered immunity in the host plant. According to these results, the authors supposed this protein to be a potential trigger of gall formation (Zhao et al., 2015).

In aphids, the salivary glands of *Hormaphis cornu* making galls on the witch-hazel (*Hamamelis virginiana*) produce a specific determinant of gall color (DGC) protein. The production of this protein is associated with the regulation of anthocyanin synthesis in the host plant (Korgaonkar et al., 2021). Anthocyanins are pigments responsible for the formation of red, purple, and blue colors in plants. In the study of Korgaonkar et al. (2021), the hyperproduction of red pigment in forming galls was correlated with the upregulation of seven genes coding enzymes acting in anthocyanin synthesis in the plant and simultaneous hyperproduction of the DGC protein in the insect. The authors supposed the triggering of red gall to be due to the injection of this potential effector protein from salivary glands into a plant tissue. Furthermore, the gene encoding DGC displays a high dN/dS ratio indicating positive selection. Thus, it shows a potential role of this gene in the galling in the context of the evolutionary arms race of aphids and their host plants.

The following studies were dedicated to another group of gall inducing organisms, gall wasps (Hymenoptera: Cynipidae). Cynipid wasps include at least 1400 species being the second largest group of gall-inducing insects after gall midges (Diptera: Cecidomyiidae), and occur on all continents, except for the Antarctic (Ronquist et al., 2015). Each gall wasp species typically attacks a single host plant species or genus. In addition, each species causes a particular form of gall in a particular plant organ. Gall forms vary from little spheres to complex multi-chamber structures (Stone and Schönrogge, 2003).

In Cynipidae a study (Gobbo et al., 2020) demonstrated positive selection for the genes associated with gall formation in *Synergus itoensis* (Cynipidae: Synergini), the only one known gall inducer species from the genus *Synergus*. The authors calculated a pairwise dN/dS ratio between *S. itoensis* and several inquiline *Synergus* species and performed gene set enrichment analysis of the genes showing a higher dN/dS. Cynipid inquiline is a form that had lost the ability to induce galls *de novo* but occupies the galls induced by a gall inducer species (Ronquist, 1994). The gene set of *S. itoensis* showing signature of positive selection, unlike those of the inquiline species, was enriched in the ‘ovarian follicle cell development’, ‘heart development’, ‘axonogenesis’, and ‘axon development’ terms (Gobbo et al., 2020). The genes enriched in these terms were supposed to reflect the ability to induce galls. The authors hypothesized that the secretions coating the egg surface are known to induce plant immunity (Dobens & Raftery, 2000; Hilker & Fatouros, 2015). The plant immune response was supposed to accompany the initial steps of gall induction just after oviposition (Gobbo et al., 2020).

Other candidate genes acting on gall formation were found in the venom glands of *Diplolepis rosae* and *Biorhiza pallida.* Transcriptome analysis revealed the overexpression of genes encoding serine proteases, phospholipases, lipases, esterases, and peroxidases (Cambier et al., 2019). These enzymes have no evident role in regulation of plant immunity or plant development. Nonetheless, in the *B. pallida* venom the authors also detected cellulases of bacterial origin, which was supposed to contribute to the lysis of a plant cell wall. Furthermore, another transcriptome study (Hearn et al., 2019) showed the overexpression of genes encoding different plant cell wall degrading enzymes (PCWDEs) like pectate lyases, rhamnogalacturonan lyases, and cellulases in *B. pallida* larvae. PCWDEs were both encoded in the insect genome and most certainly acquired via horizontal gene transfer from bacteria.

In this study, we aimed to reveal candidate genes that could be responsible for the initial stages of gall formation, i.e. gall initiation in *Diplolepis rosae*, a holarctic gall wasp causing bedeguars in wild roses *Rosa* spp. sect. Caninae (Rosaceae). *D. rosae* is a univoltine species reproducing mostly by thelytokous parthenogenesis, where virgin females produce females (Nordlander, 1973; Stille & Dävring, 1980; Heimpel & De Boer, 2008). Adults emerge from May to June and are synchronized with the development of suitable host plant tissues for gall induction (Shorthouse & Floate, 2010). After emergence, the females immediately oviposit into epidermal plant cells located between the developing leaflets of an expanding bud (Bronner, 1985). Once eggs are laid, gall tissue begins to develop. One gall can contain up to one hundred larvae (Rizzo & Massa, 2006). The feeding larvae are surrounded by gall cells and spend at the pre-nymph stage (from early November) before the next spring (Shorthouse & Floate, 2010).

Firstly, we hypothesized that gall wasps must generate new genetic variants to evade plant immune system and successfully manipulate host plant metabolism (Van Valen, 1973). Therefore, we sought to identify evidence of positive selection in the genome of *D. rosae* by performing a test for selection as in the study of Korgaonkar et al. 2021. Secondly, we employed transcriptomics and measured gene expression levels in various *D. rosae* tissues at different steps of gall formation. Our aim was to identify genes that are overexpressed during the early stages compared to the later stages. Subsequently, we expected to refine the list of candidate genes by comparing two sets of candidates identified by both approaches. On the one hand, we sought to exclude genes that might be under selection but not be expressed during the initial stages of gall formation. On the other hand, the set of genes under positive selection could help to exclude conserved genes overexpressed during gall initiation but likely responsible for other processes, such as insect development.

## Results

### Relative differential gene expression data analysis

BRAKER annotation (.*gff3* file) (Stanke et al., 2006, 2008; Hoff et al., 2016, 2019; Bruna et al., 2021) of the *D. rosae* revealed 125,626 genes, 135,510 mRNA transcripts, 346,652 exons and protein-coding sequences (CDS), and 211,236 introns. In the RNAseq analysis, the total number of reads aligned to the *gff3.* annotation varied between 98.61 million and 151.2 million depending on the library. The percentage of reads overlapping with the *D. rosae* genes varied between 38.72 % and 69.45 % (**Table S1**). Pairwise differential expression analysis revealed genes specifically expressed in the *D. rosae* egg (1028 genes), mid-July larva (2390), early September larva (455), October salivary gland (112), November salivary gland (2516), and female adult head (855) (**Figure 1, Table S2**). The number of genes upregulated during the whole process of gall formation (combined sample ‘mid-July *D. rosae* larva + mid-July *D. eglanteriae* larva + early September *D. rosae* larva + October *D. rosae* salivary gland + November *D. rosae* salivary gland’ vs combined sample ‘head + egg’) was 11,916. Genes encoding proteins containing leucine-rich repeat, plant cell wall degrading enzymes, and venom-like enzymes were found to be up-regulated during the whole process of galling.

**Figure 1.**
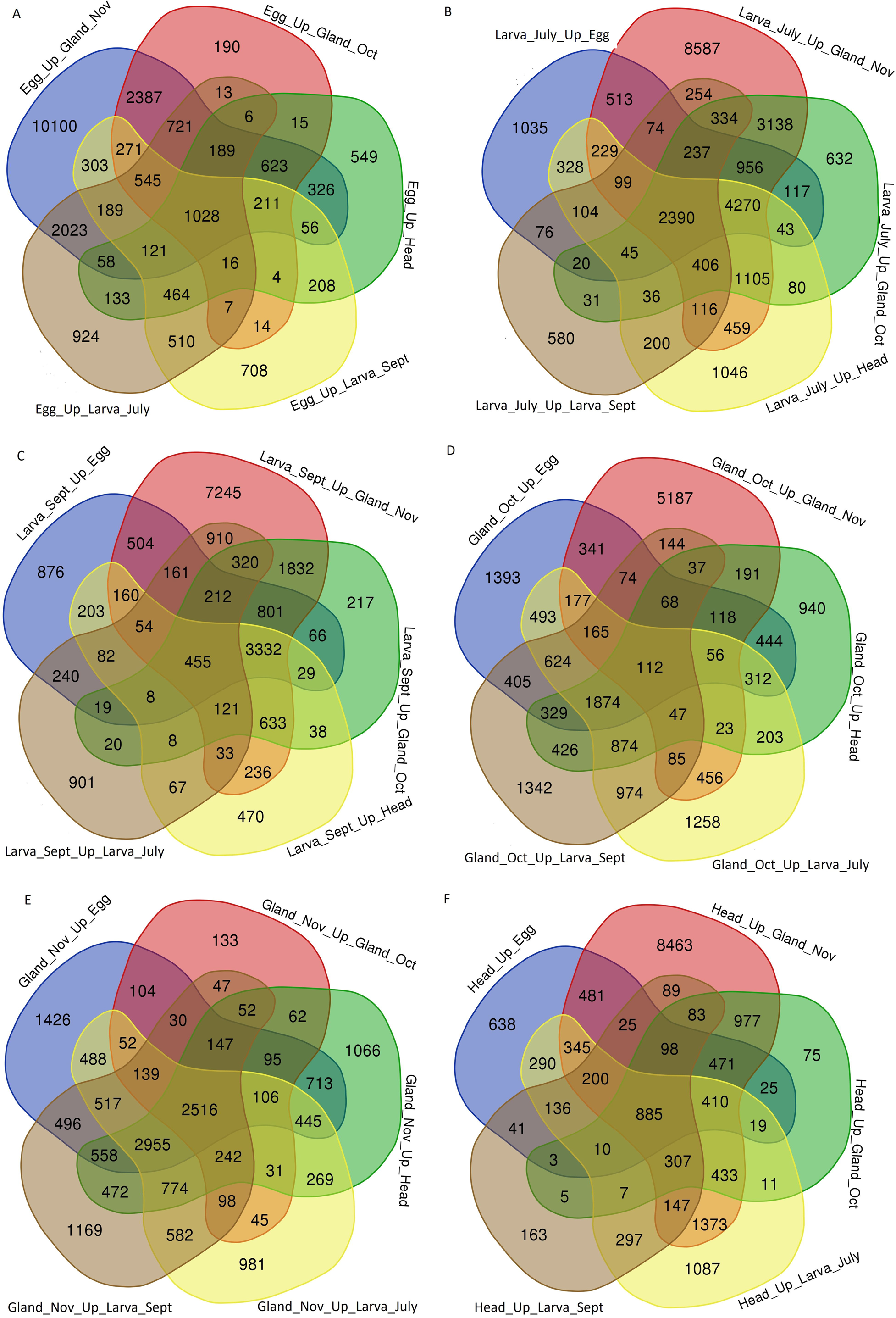

### Gene annotations and dynamics of gene expression

The total number of annotated genes by eggNOG (Cantalapiedra et al., 2021; Huerta-Cepas et al., 2021) was 62,690 including 47,871 genes annotated in Cluster of Orthologous Groups (COG) database and 12,461 genes in the Gene Ontology (GO). Among 11,916 genes up-regulated at least at one stage during the whole process of gall formation (combined sample ‘mid-July *D. rosae* larva + mid-July *D. eglanteriae* larva + early September *D. rosae* larva + October *D. rosae* salivary gland + November *D. rosae* salivary gland’ vs combined sample ‘female adult head + egg’), 5,152 genes were annotated at the protein families level (Finn et al., 2016), 4,088 genes were annotated with COG, and 2,208 genes were annotated with GO. The number of up-regulated genes encoding proteins containing leucine-rich repeats (LRR), plant cell wall degrading enzymes (PCWDE), and venom-like enzymes was 15, 6, and 30, respectively (**Table S3**). Five genes encoding LRR proteins and 13 genes encoding venom-like enzymes showed greater expression during July and September (active gall growth) compared to October and November (**Figures 2-4**). Among PCWDEs, the gene g94279 encoding cellulase was highly expressed from July to November (**Figure 5**). The other genes encoding PCWDEs showed higher expression from July to October, followed by a decline in November.

**Figure 2.**
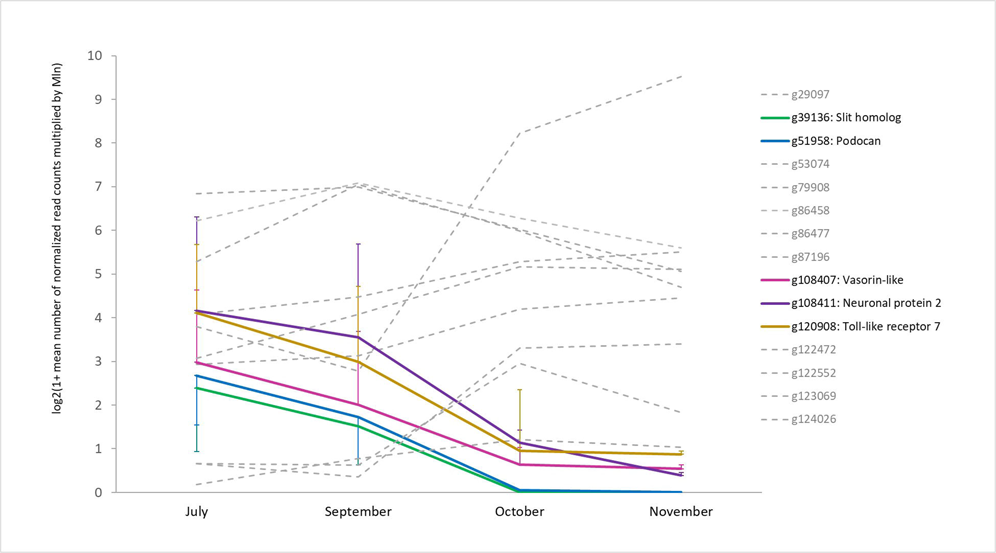

**Figure 3.**
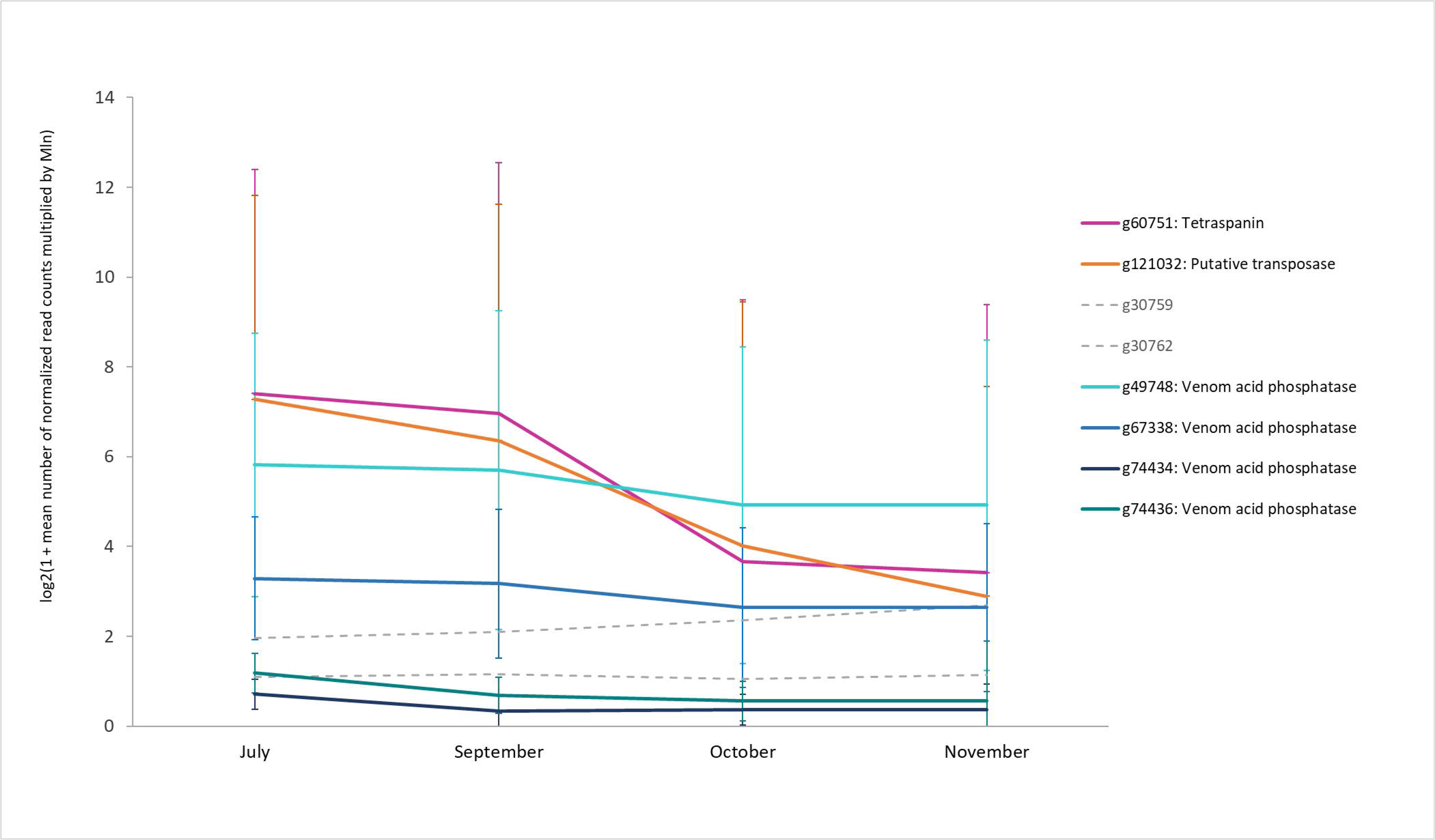

**Figure 4.**
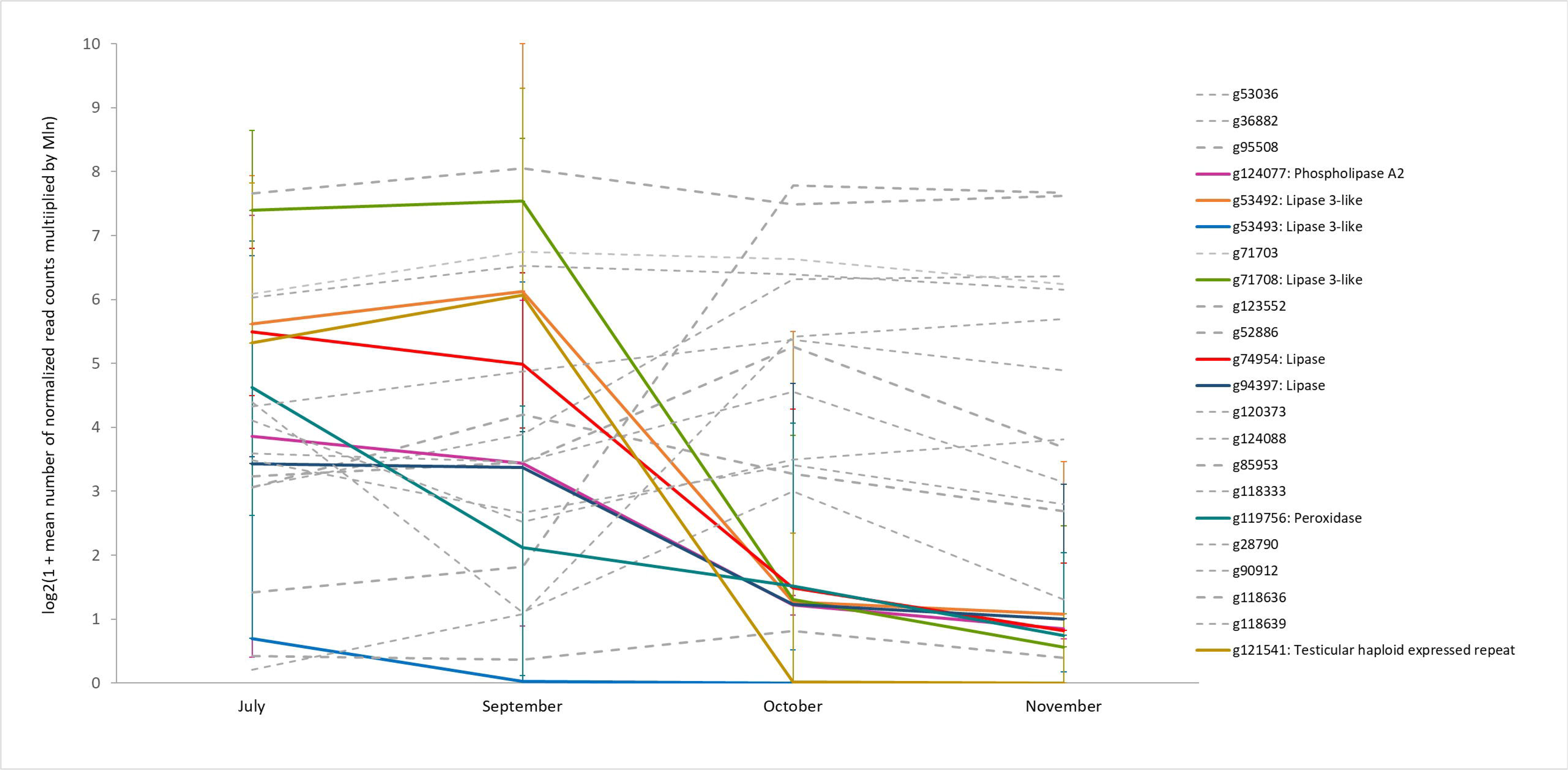

**Figure 5.**
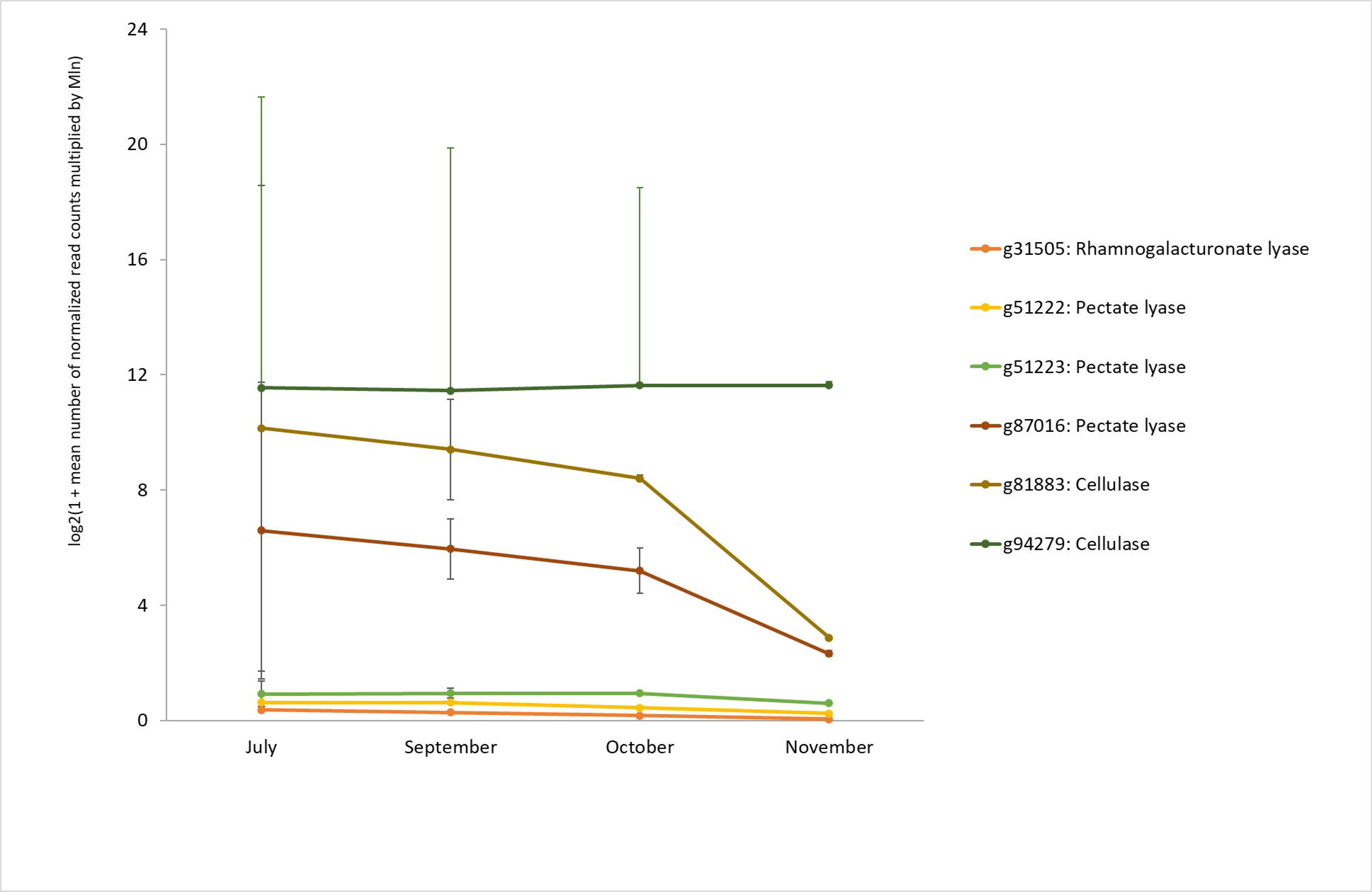

### McDonald–Kreitman test

The initial number of extracted alignments (concatenated protein-coding sequences, CDS) was 135,472. The number of alignments showing at least one polymorphic or divergent site was 120,442. The number of alignments showing at least one divergent site was 266. Genome-wide selection coefficient *s* was estimated at −2.1. Among genes specifically expressed in the mid-July larva, the early September larva, during active gall growing (from July to September), and during whole gall formation (from July to November), one gene was under positive selection (mean *s* estimation provided by SniPRE (Eilertson et al., 2012) was 0.74), 348 genes were under neutral selection (mean *s*: [-1.49; 0.38]), and 11,567 genes were under negative selection (mean *s*: [−4.32; −0.60]) (**Figure 6**). The gene (g50314) under positive selection encoded a transposable element.

**Figure 6.**
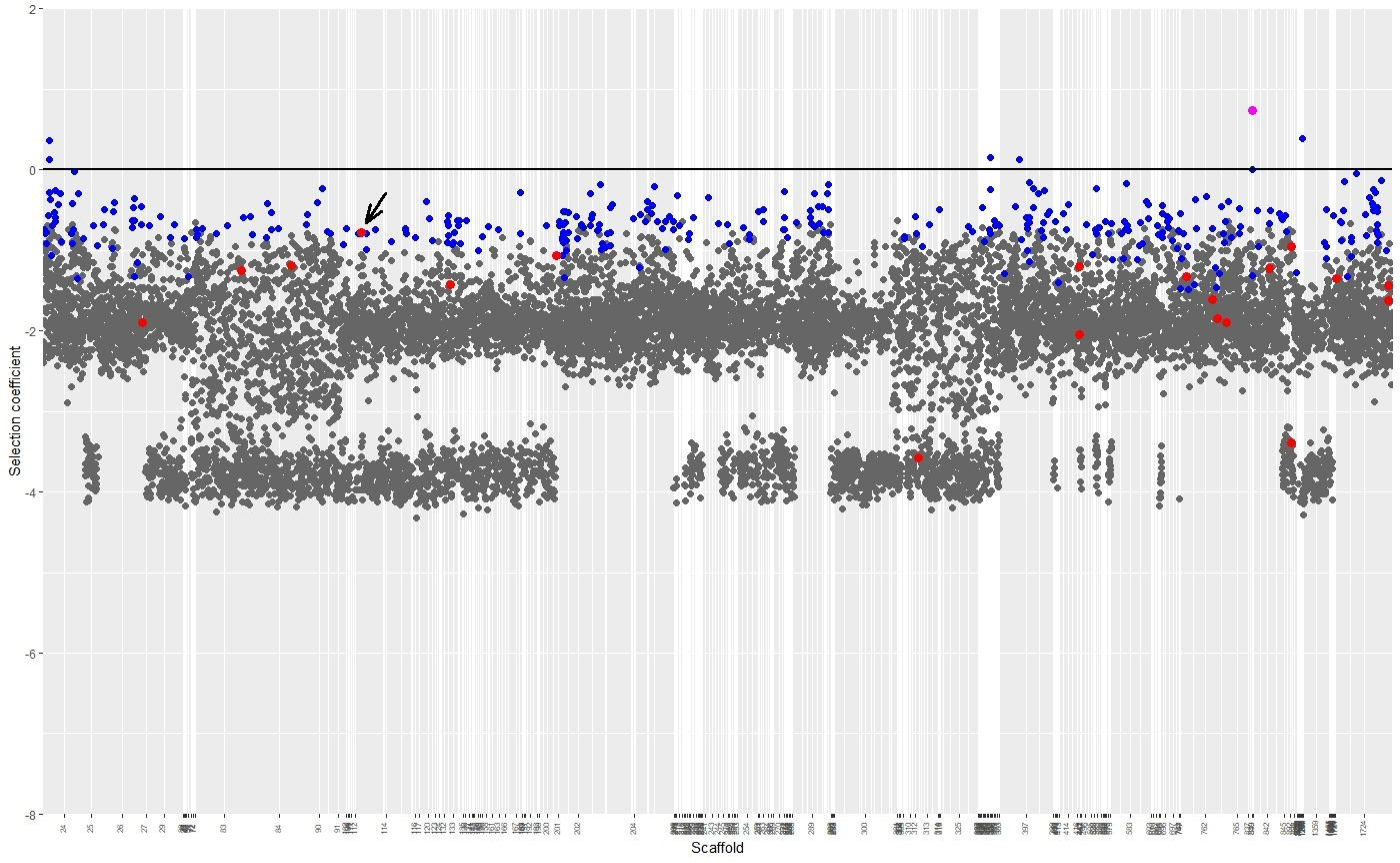

## Discussion

In this study, we performed the transcriptome analysis of *D. rosae* to identify candidate genes involved in gall formation. We focused on the genes that were up-regulated during the earliest observable *D. rosae* larval stages in nature, i.e. mid-July and early September. We gave a particular focus on the candidate genes that were similarly annotated as those previously reported in other studies of gall-inducing insects (Zhao et al., 2015; Cambier et al., 2019; Hearn et al., 2019) for two reasons. The first reason is the absence of a significant number of gene annotations. Among the genes that were up-regulated at least at one stage from July to November, only 43% of genes had available protein family annotations. Orthologous Group and Gene Ontology annotations were available for 34% and 19% of the up-regulated genes, respectively. Hence, performing an enrichment analysis solely on genes with available functional annotations could lead to inaccurate results. The second reason is that approximately half of the available functional annotations were attributed to transposable elements, whose role in galling is challenging to evaluate; the simultaneous overexpression of genes encoding transposable elements can be explained by an active transcription occurring during insect development. Thus, we investigated the *D. rosae* genes (1) encoding any proteins with the leucine-rich domain, as detected in the Hessian fly salivary gland protein (Zhao et al., 2015), (2) similar to those overexpressed in the *D. rosae* venom gland (Cambier el al., 2019), and (3) orthologous to those overexpressed in the *B. pallida* larva and venom gland (Cambier el al., 2019; Hearn et al., 2019).

The first group of up-regulated genes in the mid-July and early September *D. rosae* larvae encodes leucine-rich repeat (LRR) proteins: slit homolog, neuronal protein 2, podocan, vasorin-like protein, and Toll-like receptor 7. Slit homolog is involved in neural development and controls axon crossing in *Drosophila* (Brose et al., 1999; Kidd et al., 1999). The peptide matching the LRR domain of neuronal protein 2 is likely to be involved in synapse functioning (Linhoff et al., 2009). Thus, the expression of these genes can be related to insect development rather than being involved in galling. Other genes encoding LRR proteins can be associated with insect immune response. For example, podocan homolog was shown to be overexpressed in *Cnaphalocrocis medinalis* (Lepidoptera: Crambidae) in response to a baculovirus infection (Han et al., 2021). Vasorin-like protein and Toll-7 receptor belong to Toll-like receptors, a group of transmembrane proteins containing extracellular LRR motifs that play an essential role in invertebrate immunity and contribute to embryonic development in insects. These receptors recognize specific molecular patterns of various pathogenic microorganisms and initiate immune response (Medzhitov, 2001; Leulier & Lemaitre, 2008). Notably, Fjøsne et al. (2015) showed the vasorin-like protein to be overexpressed in *Dentrobaena veneta* (Annelida: Lambricidae) in response to a bacterial infection. Another study (Park et al., 2019) demonstrated that the expression of Toll-like receptor 7 was induced by both bacterial and fungal infections in *Tenebrio molitor* (Coleoptera: Tenebrionidae).

The second group of examined genes was those encoding proteins previously detected in the *D. rosae* venom gland (Cambier et al., 2019): tetraspanin and putative transposase. Additionally, we examined the genes annotated as ‘venom acid phosphatase’ and orthologous to those up-regulated in the *B. pallida* venom gland (Cambier et al., 2019). Tetraspanins are a group of proteins implicated in multiple biological processes including insect development, reproductive processes, extracellular matrix organization, vesicle formation, and host-pathogen interactions (Todres et al., 2000; Hemler, 2003). In host-pathogen interactions, tetraspanin serves as surface marker of immune cells and involved in signal transduction when generating immune response (Zhuang et al., 2007). For instance, Mei et al. (2023) showed overexpression of tetraspanin when *Bombyx mori* (Lepidoptera: Bombycidae) was exposed to a viral infection, and Mahadav et al. (2008) demonstrated that the parasitism by the wasp *Eretmocerus mundus* (Hymenoptera: Aphelinidae) induced the expression of tetraspanin in *Bemisia tabaci* (Hemiptera: Aleyrodidae). Other genes overexpressed in both *D. rosae* venom glands (Cambier et al., 2019) and the young larvae encode venom acid phosphatases, phospholipases A2, lipases, and peroxidase. Notably, some of these enzymes were detected in venoms of various parasitic Hymenoptera (Colinet et al., 2013; Poirié et al., 2014). The main role of parasitoid venom is to inhibit the insect host immune response. Common examples of immunomodulating parasitoid venom components are serine proteases (Asgari et al., 2003) and serine protease inhibitors (Colinet et al., 2009; Qian et al., 2015) that interrupt the formation of the melanin protective capsule in the insect host. However, the function of the other hydrolytic enzymes in regulation of the host immunity is unclear or has not been confirmed (Dani et al., 2005; Colinet et al., 2013; Dorémus et al., 2013; Poirie et al., 2014). Hence, the uncertain role of these proteins in parasitoid venoms presents a challenge when hypothesizing their role in gall wasps. Nonetheless, we can suppose that the hyperproduction of such proteins is due to insect development and immune challenges in the young *D. rosae* larvae. Firstly, lipases and phospholipases A2 are likely to be implicated in fatty acid metabolism during the gall wasp development. This could find support in the study of Akpinar et al. (2017) that demonstrated changes in fatty acid composition during the whole life cycle of *Diplolepis fructuum*. Secondly, these enzymes are involved in various processes including lipid metabolism and lipid signaling in insect development, reproduction, neurotransmission, and immune responses (Stanley, 2006; Shrestha et al., 2010; Hossen et al., 2016). Another enzyme showing diverse functions in insects is acid phosphatase playing a role in different biosynthetic and signaling pathways (Xia et al., 2000; Hossen et al., 2016; Lehmann, 2021). An increase in peroxidase activity might be necessary to deactivate toxic molecules and radicals generated during the immune response that could take place during the gall wasp development. Indeed, *D. rosae* larvae are exposed to high parasitic pressure: the percentage of parasitoid individuals in the *D. rosae* gall community can reach more than 70% (Rizzo & Massa, 2006; Todorov et al., 2012). Furthermore, one of the most common parasitoids of *D. rosae*, *Orthopelma mediator* (Hymenoptera: Ichneumonidae), attacks *D. rosae* even before the gall begins to develop (Stille, 1984). This can trigger an immune response in the gall wasp larvae at the early stages of gall formation. In addition, *D. rosae* can be also exposed to endophytic microorganisms. For instance, the microbiome of *Rosa* spp. consists of bacteria, such as *Bacillus* and *Staphylococcus* (Xia et al., 2020), as well as fungi (Zhao et al., 2018). Furthermore, hyperproduction of peroxidase in the young *D. rosae* larvae may be regarded as a response to the defensive mechanisms of the host plant. When attacked by herbivore organisms, plants initiate a defensive cascade, releasing signaling molecules such as ATP, Ca^2+^, and H_2_O_2_. These molecules trigger the expression of defense genes and, subsequently, plant defense responses (Guiguet et al., 2016). Hence, peroxidase of *D. rosae* could neutralize H_2_O_2_ produced by the host plant, thereby allowing it to evade plant immune response. Furthermore, the hyperproduction of enzymes like peroxiredoxin, which are also involved in the breakdown of H_2_O_2_, was detected in salivary glands of various insect herbivores (Guiguet et al., 2016). Lastly, in summer *D. rosae* larvae, we detected the overexpression of a gene encoding testicular haploid expressed repeat, which may be involved in the development of the insect’s reproductive system.

The third group comprises *D. rosae* genes that are orthologous to those encoding plant cell wall degrading enzymes (PCWDE) in *B. pallida* (Hearn et al., 2019): cellulase, pectate lyase, and rhamnogalacturonate lyase. Similar to other insect herbivores, the *D. rosae* PCWDE genes were acquired from bacteria via horizontal gene transfer (Wybouw et al., 2016). The PCWDE genes are found in cynipid gallers and inquilines, which is associated with their role in the formation of gall chambers for developing larvae. Furthermore, the PCWDEs genes have been lost in the members from the nested parasitoid family Figitidae, or are presented as fragments of functional domains or pseudogenes (Hearn et al., 2019, 2023). In *D. rosae*, we observed that the PCWDE genes showed the same expression level during the whole examined period of galling, i.e. July-November. However, we could suppose that PCWDEs may play a role not only in the formation of a gall chamber during larval nutrition but also in the earlier steps of gall formation, i.e. initiation. Indeed, the functioning of these enzymes leads to the release of degradation products serving as plant signaling molecules (Vallarino & Osorio, 2012; De Lorenzo et al., 2019; Hearn et al., 2019). These metabolites could potentially modulate the differentiation of plant cells and provoke the development of gall tissue. Furthermore, the enzyme such as cellulase was detected in the *B. pallida* venom gland (Cambier et al., 2019). It could contribute to the degradation of plant cell wall during oviposition and also lead to release of metabolites, thereby initiating gall formation. Regrettably, due to the limitations in data quality (Cambier et al., 2019), we do not dispose of complete information about the venom composition of *D. rosae* that could provide an additional argument. Nevertheless, *D. rosae* remains another example of Cynipidae highlighting the role of the genes encoding PCWDEs in gall formation, particularly during the growth stage.

As we have observed, it is challenging to link the studied genes with gall formation in *D. rosae*. Most of the examined genes can be predominantly involved in insect development and nutrition rather than host plant manipulation. Nonetheless, we observed that various up-regulated genes were associated with the immune response of insects. The last observation highlights that almost all genes were under negative selection, and only one gene encoding a transposable element was under positive selection. It can be explained by the asexual mode of reproduction in *D. rosae*, i.e. thelytokous parthenogenesis via gamete duplication (Nordlander, 1973; Stille & Dävring, 1980; Heimpel & De Boer 2008; Mozhaitseva et al., 2023). This mode of reproduction results in complete homozygosity where all alleles are in complete linkage. Consequently, in highly homozygous thelytokous *D. rosae* females, disadvantageous alleles and all linked alleles are purged quickly (Pearcy et al., 2006). However, positive selection may occur but remain undetectable due to the extremely low recombination rate in *D. rosae* (Mozhaitseva et al., 2023). Males constitute only 0 - 4% of *D. rosae* populations (Nordlander, 1973; laboratory observations) and have minimal contribution to each generation. This leads to the phenomenon of clonal interference (Muller, 1932). In asexually reproducing organisms, a beneficial mutation takes a longer time to be fixed or can be lost in the absence of recombination and the inability to spread in a population. Thus, we were unable to detect the signatures of positive selection such as elevated relative frequency of a non-synonymous polymorphism in the protein-coding sequences of *D. rosae*.

## Conclusion

Transcriptome analysis revealed the overexpression of 11,916 *D. rosae* genes during the observable stages of gall formation (mid-July to early November), with 7,153 genes being overexpressed specifically during the active gall growth phase (mid-July to early September). Among the examined genes, those encoding plant cell wall degrading enzymes can be associated with galling. The other upregulated genes are likely to be implicated in insect development and the immune response to parasitoids and the host plant microbiota. Almost all genes have been found to be under negative selection. It could be explained by the mostly asexual mode of reproduction in *D. rosae*, which results in a rapid purge of deleterious and all linked alleles. Besides, positive selection could be undetectable because of reduced recombination in this species and the effect of clonal interference.

Further investigations could be focused on designing experiments that allow to examine the *D. rosae* transcriptome sampled during gall initiation. One should master the life cycle of *D. rosae* in greenhouse conditions. It could help to pinpoint the moment and location of oviposition during the experiment and measure gene expression levels in *D. rosae* eggs and young larvae after the first hours and days. Additionally, other groups of genes, such as those involved in the disrupting danger signal molecules in plants, could be examined using the same methodology as in this study. Improving gene annotations and conducting enrichment analyses would help to identify gene groups with specific functions. Lastly, we could analyze the evolution of candidate genes associated with galling within Cynipidae s. lat. and infer whether any genes present in gall inducers and absent in their parasitic members.

## Experimental procedures

### Sampling

Seventeen *D. rosae* and two *D. eglanteriae* galls were collected in France from September 2019 to December 2020 (**Table S4**). *D. rosae* individuals were collected from the same host plant specimen and came from the one female due to a clonal mode of reproduction (Mozhaitseva et al., 2023). After sampling, the galls were kept in plastic bags at room temperature. Insect material was removed by dissection of gall tissue for further DNA extraction.

### DNA extraction for Illumina sequencing

The larvae were homogenized in 2-mL plastic tubes by a TissueLyser (Qiagen) with adding a metallic bead. The initial mass of larvae varied between 1.8 and 56.0 mg. DNA was extracted using a DNeasy Blood and Tissue Kit (QIAGEN, Germany). 260/280 and 260/230 ratios varied from 1.35 to 2.08 and from 0.40 to 2.08, respectively. DNA quantity ranged from 0.42 to 6.9 μg. Before the sequencing, *D. eglanteriae* was verified using the PCR of the gene encoding cytochrome oxidase c subunit I to distinguish between the gall wasp larvae and their parasitoids. The sequence of a PCR product (**Figure S1**) was verified in the Nucleotide collection (nt) database (the NCBI platform: https://blast.ncbi.nlm.nih.gov). After that, the Illumina sequencing was performed by Genotoul, Toulouse, France (https://get.genotoul.fr).

### SNP calling

All reads of *Diplolepis spp.* were firstly aligned to the *D. rosae* reference (accession JAPYXD000000000 in DDBJ/ENA/GenBank) using Bowtie 2 v. 2.3.5.1 (Langmead & Salzberg, 2012) with --end-to-end (default, *D. rosae* reads) or --local (*D. eglanteriae* reads) mode. The alignments were processed for further manipulations using Samtools 1.7 (view, sort, index, depth, and stat options) (Danecek et al., 2021). The quality of each alignment was estimated by calculating the average percentage of the mapped reads (56.6% - 98.1%), the average quality (35.4 - 35.7), the average coverage (22x - 119x), and the error rate (0.0079 - 0.015). After that, the identification of polymorphisms across the genome was performed by using a pipeline proposed by GenomeAnalysisToolkit (GATK) v. 4.0 (AddOrReplaceReadGroups, HaplotypeCaller, CombineGVCFs, and GenotypeGVCFs tools) (https://gatk.broadinstitute.org/hc/en-us/articles/360035890411-Calling-variants-on-cohorts-of-samples-using-the-HaplotypeCaller-in-GVCF-mode). Indels and non-biallelic sites were removed from the *.vcf* file using bcftools v. 1.7 (filter command) (Danecek et al., 2021). Phasing of the *.vcf* file was performed by beagle v. 5.4 (Browning et al., 2021) to show two haplotypes corresponding to each individual (**Code S1**).

### McDonald–Kreitman test

The *.fasta* alignments were extracted from the phased *.vcf* file using the vcf2fasta.py program (https://github.com/santiagosnchez/vcf2fasta). Gene coordinates were shown in a *.gff* annotation file and the reference *.fasta* file. Each alignment corresponded to concatenated coding sequences (CDS) belonging to the same transcript. For each *D. rosae* alignment, the number of polymorphic synonymous/non-synonymous and divergent synonymous/non-synonymous mutations was inferred by the MKtest program from libsequence (Thornton, 2003) using *D. eglanteriae* as an outgroup. The mean number of synonymous and replacement sites per gene was calculated by the polydNdS program from libsequence (Thornton, 2003). Obtained data were used to estimate the direction of selection, the mean value of selection coefficient *s* and lower and upper *s* bounds using SnIPRE, a McDonald-Kreitman type analysis based on a generalized linear mixed model (Eilertson et al., 2012) (**Code S2**).

### RNA extraction

*D. rosae* galls developing on the same dog rose were sampled from May 2022 to November 2022 in Bures-sur-Yvette (48°42′12″N, 2°9′35″E). *D. eglanteriae* dog rose galls were sampled in mid-July 2022. The insect material was removed by gall dissection. Before RNA extraction, the insects were kept in a 2-ml Eppendorf tube containing 1 ml of RNAlater Solution (Invitrogen, Lithuania) at 4 °C. Insect tissue was homogenized by TissueLyser (Qiagen) using a metallic bead. RNA was extracted using a RNeasy Mini Kit (QIAGEN, Germany). After extraction, the purity of the samples was estimated using 260/280 (varied between 1.90 and 2.25) and 260/230 (varied between 0.31 and 2.25) ratios measured by a Thermo Scientific NanoDrop 2000 Spectrophotometer. RNA quantity was estimated using a Qubit RNA BR Assay Kit (Molecular Probes, USA) and a Qubit Fluorometer and varied from 0.82 to 5.3 µg.

### Sampling

Since we did not master the life cycle of *D. rosae* for a laboratory experiment, we collected the galls in natural conditions. We obtained the following samples:

- mid-July larva (first visible gall);
- early September larva (growing gall);
- early October larva salivary glands (mature gall);
- early November larva/pre-nymph salivary glands (wintering gall);
- head from the female adult emerging from the gall kept in the laboratory (control sample);
- egg removed from the emerged female adult (control sample);
- additionally, mid-July *D. eglanteriae* larva to exclude species-specific genes.

We expected to detect candidate genes specifically expressed:

- at each life cycle stage or tissue: for instance, to show up-regulated genes in the mid-July *D. rosae* larva, we searched for an overlap between the up-regulated genes by comparing the pairs:

`o` mid-July larva - egg,
`o` mid-July larva - early September larva,
`o` mid-July larva - early October larva salivary glands,
`o` mid-July larva - early November larva salivary glands,
`o` mid-July larva - female adult head;
- during the active gall growth (’growth’ vs no ‘growth’): comparison between the sample set pair ‘mid-July larva + early September larva + mid-July *D. eglanteriae* larva’ and ‘egg + early October larva salivary glands + early November larva salivary glands + female adult head’,
- during the whole gall formation (‘gall’ vs ‘no gall’): comparison between the sample set pair ‘mid-July larva + early September larva + mid-July *D. eglanteriae* larva + early October larva salivary glands + early November larva salivary glands’ and ‘egg + female adult head’.

### Relative differential gene expression data analysis

cDNA sequencing and library preparation (Illumina NovaSeq 50 M 150-bp reads, PolyA enrichment, non-stranded)) was performed by Novogene Europe (Cambridge, UK). Quality control of raw reads was performed using the fastQC program v. 0.12.1 (Andrews, 2010) (**Table S5**). Then, the reads were aligned to the genome reference by STAR v. 2.7.10b (Dobin et al., 2013). Genome coordinates were provided in a *.gff3* file generated by BRAKER v. 3.0.3 (Stanke et al., 2006, 2008; Hoff et al., 2016, 2019; Bruna et al., 2021) from the reference genome and .bam alignments obtained at the previous step. The count of aligned reads to predicted genes was performed by featureCounts v. 2.0.6 (Liao et al., 2014) using gene coordinates given in the *.gff3* gene prediction file. The counting was performed at gene level. The summarizing gene count matrix was then used in the relative differential expression analysis performed by DESeq2 v. 1.36.0 (Love et al., 2014). Up- and down-regulated genes specifically expressed at each stage or tissue were assessed by pairwise comparison of the following samples: egg, mid-July larva, early September larva, October salivary glands, November salivary glands, and female adult head. Data quality was assessed by calculating pairwise Euclidean distances (**Figures S2-S4**), performing the principal component analysis (**Figures S5-S7**), and plotting dispersion estimates (**Figures S8-S10**).

### Gene annotation and alignment

*D. rosae* genes were functionally annotated by eggNOG-mapper v. 2.1.7 (Cantalapiedra et al., 2021; Huerta-Cepas et al., 2021) and using the Non-redundant protein sequences (nr) database (the NCBI platform).

Firstly, we examined whether the genes specific to the *D. rosae* venom gland (Cambier et al. 2019) were upregulated during the early stages of gall formation. The reads from one available adult female venom gland sample (run SRR8501630) and one available adult female ovary sample (run SRR8501629) obtained from the SRA database (the NCBI platform) were aligned to the reference genome by STAR v. 2.7.10b (Dobin et al., 2013). The count of reads aligned to the annotated genes was performed by featureCounts v. 2.0.6 (Liao et al., 2014). The number of read counts was then normalized by the total number of reads in the respective library. According to Cambier et al. (2019), genes showing at least 20-times higher number of read counts in the *D. rosae* venom gland compared to the ovary were considered up-regulated. Finally, the presence of these genes was assessed in the list of those overexpressed in the mid-July and the early September *D. rosae* larvae by aligning the gene sets using blastn of BLAST v. 2.14.0 (Camacho et al. 2009).

Secondly, we examined whether *D. rosae* orthologous genes encoding venom components (Cambier et al., 2019) and plant cell wall degrading enzymes (Hearn et al., 2019) in *B. pallida* were overexpressed during gall formation. Candidate orthologous genes overexpressed in the mid-July *D. rosae* larva, the early September larva, October salivary glands, and November salivary glands were identified by applying the bidirectional best hit strategy (Smith & Waterman, 1981). First, the *D. rosae* genes (protein sequence, -query tag) were aligned to the *B. pallida* transcriptome (-db) (Hearn et al. 2023) by tblastn of BLAST v. 2.14.0 (Camacho et al. 2009). Next, the *B. pallida* sequences (-query) showing the highest bit score were aligned to the *D. rosae* genes (-db) by blastx of BLAST v. 2.14.0 (Camacho et al. 2009). The -db *D. rosae* genes showing the highest bit score were then compared with the -query *D. rosae* genes from the tblastn output: if the same *D. rosae* gene matched the same *B. pallida* sequence, they were considered orthologs.

### Statistics

All statistical analyses were performed by R v 4.2.2 (R Core Team, 2022). The significance level was set to 0.05.

## Author Contributions

**Ksenia Mozhaitseva:** writing – original draft; writing – review and editing; sample collection; methodology; data analysis; visualization. **Zoe Tourrain:** writing – review and editing; sample collection, methodology, visualization. **Antoine Branca:** writing – review and editing; sample collection; methodology; supervision; funding acquisition; project administration.

## Acknowledgements

We would like to thank our colleagues from EGCE and ESE laboratories for fruitful discussions that helped to enrich the quality and depth of this study.

## Data Availability Statement

The reference genome of *D*. *rosae* is available at the NCBI platform (https://www.ncbi.nlm.nih.gov/): a DDBJ/ENA/GenBank accession is JAPYXD000000000 (version JAPYXD000000000.1), a BioProject accession is PRJNA914909, and a BioSample identifier is SAMN32363506. *D*. *rosae* Illumina reads are available at the NCBI platform: SRA accessions are SRS16596603−SRS16596619, and BioSample accessions are SAMN32903234−SAMN32903250. *D. eglanteriae* reads are available under accessions *M*−*N*. Illumina reads for RNAseq are available under accessions *X−Y*.

