## Supplementary Material for "Transcriptome analysis of *Diplolepis rosae*: revealing overexpression of genes potentially associated with insect immune response and gall formation at early larval stages"

**Tables**

**Table S1.** Percentage of different classes of reads aligned to the annotated genes of *Diplolepis rosae*.

| **Sample** | **% Assigned** | **% Unassigned_**  **NoFeatures** | **% Unassigned_**  **Ambiguity** | **Total number of reads** |
| --- | --- | --- | --- | --- |
| Egg_1 | 46.57 | 52.40 | 1.03 | 105,692,304 |
| Egg_3 | 50.60 | 48.40 | 1.00 | 109,586,716 |
| Egg_4 | 53.18 | 45.81 | 1.01 | 99,616,178 |
| Gland_Nov_1 | 66.81 | 31.92 | 1.27 | 128,332,522 |
| Gland_Nov_2 | 66.20 | 32.46 | 1.34 | 141,362,650 |
| Gland_Nov_3 | 69.45 | 29.06 | 1.49 | 151,175,994 |
| Gland_Nov_4 | 68.03 | 30.54 | 1.43 | 117,059,536 |
| Gland_Oct_1 | 53.06 | 45.92 | 1.02 | 115,042,208 |
| Gland_Oct_2 | 38.72 | 60.47 | 0.81 | 137,425,368 |
| Gland_Oct_3 | 64.30 | 34.44 | 1.26 | 119,692,118 |
| Gland_Oct_4 | 59.03 | 39.80 | 1.17 | 118,450,874 |
| Head_1 | 41.52 | 57.64 | 0.84 | 122,053,964 |
| Head_2 | 43.12 | 56.02 | 0.86 | 120,933,152 |
| Larva_July_1 | 61.56 | 36.68 | 1.76 | 98,612,530 |
| Larva_July_2 | 56.60 | 41.80 | 1.60 | 101,374,936 |
| Larva_July_3 | 63.17 | 35.61 | 1.22 | 105,756,354 |
| Larva_July_4 | 65.36 | 32.47 | 2.17 | 103,313,482 |
| Larva_Sept_1 | 50.27 | 48.57 | 1.16 | 134,714,808 |
| Larva_Sept_2 | 49.08 | 49.78 | 1.14 | 122,851,446 |
| Larva_Sept_3 | 49.95 | 48.60 | 1.45 | 115,339,990 |
| Larva_Sept_4 | 51.67 | 47.02 | 1.31 | 127,945,160 |
| D_eglant_2 | 56.10 | 24.16 | 1.25 | 151,090,542 |
| D_eglant_4 | 50.84 | 26.23 | 1.08 | 157,615,770 |

Egg: *Diplolepis rosae* eggs removed from the female adults. Gland_Nov: salivary glands from the *D. rosae* larva collected in November. Gland_Oct: salivary glands from the *D. rosae* larva collected in October. Head: *D. rosae* female adult head. Larva_July: *D. rosae* early larva removed from the gall in mid-July. Larva_Sept: *D. rosae* larva removed from the gall in early September. D_eglant: *D. eglanteriae.* Assigned: reads successfully assigned to one or another feature (gene). Only this class of reads was taken into account in the differential expression analysis. Unassigned_NoFeatures: alignments that do not overlap any feature (gene). Unassigned_Ambiguity: alignments that overlap more than one feature (gene). In the categories ‘Unassigned_Unmapped’ (unmapped reads cannot be assigned), ‘Unassigned_Read_Type’ (reads showing unexpected, i.e. no-pair-end type), ‘Unassigned_Singleton’ (read pairs showing only one read mapped), ‘Unassigned_MappingQuality’ (reads showing the mapping quality score less than the threshold of 0 by default), ‘Unassigned_Chimera’ (two ends in a paired-end alignment are located on different chromosomes), ‘Unassigned_FragmentLength’ (alignments showing the length out of the min and max thresholds, i.e. 50 bp and 600 bp by default), ‘Unassigned_Duplicate’ (duplicated reads), ‘Unassigned_MultiMapping’ (reads mapping different genome positions), ‘Unassigned_Secondary’ (secondary alignments, i.e. that can match another genome location but with a less probability than the first one), ‘Unassigned_NonSplit’ (useful if the count of exon-spanning reads is required), ‘Unassigned_Overlapping_Length’ (minimum number of overlapping bases in the alignment required to be assigned, 1 by default), the number of alignments is 0 in all samples (Liao et al., 2014).

**Table S2.** Number of differentially expressed genes after performing pairwise comparisons between different stages and tissues of *Diplolepis rosae*.

| **Pair of samples / Counts** | **Total non-zero** | **Up** | **Down** | **Outliers** | **Low counts**  **(mean count <)** |
| --- | --- | --- | --- | --- | --- |
| Larva_July vs Egg | 101,399 | 10,536 | 6,947 | 21 | 40,765 (< 3) |
| Larva_Sept vs Egg | 91,907 | 7,202 | 4,655 | 723 | 36,697 (< 3) |
| Gland_Oct vs Egg | 98,161 | 6,985 | 6,240 | 839 | 39,377 (< 3) |
| Gland_Nov vs Egg | 95,520 | 10,787 | 19,151 | 174 | 32,782 (< 2) |
| Head vs Egg | 90,432 | 4,077 | 4,007 | 2 | 41,215 (< 4) |
| Larva_Sept vs Larva_July | 100,904 | 3,611 | 5,002 | 1,234 | 44,415 (< 3) |
| Larva_July vs Gland_Oct | 105,466 | 13,840 | 7,733 | 1,293 | 38,441 (< 2) |
| Larva_July vs Gland_Nov | 103,097 | 23,167 | 10,240 | 166 | 33,589 (< 2) |
| Larva_July vs Head | 101,299 | 10,956 | 5,957 | 33 | 46,539 (< 3) |
| Larva_Sept vs Gland_Oct | 93,784 | 8,111 | 7,580 | 2,256 | 35,688 (< 3) |
| Larva_Sept vs Gland_Nov | 90,040 | 17,009 | 10,794 | 1486 | 27,337 (< 2) |
| Larva_Sept vs Head | 84,961 | 5,929 | 2,496 | 243 | 40,143 (< 5) |
| Gland_Oct vs Gland_Nov | 92,754 | 7,281 | 3,899 | 2,097 | 38,824 (< 3) |
| Head vs Gland_Oct | 92,287 | 3,819 | 6,054 | 1,089 | 42,125 (< 4) |
| Head vs Gland_Nov | 88,876 | 14,787 | 10,503 | 10 | 33712 (< 2) |
| Growth vs no_Growth | 114,798 | 11,050 | 5,293 | 0 | 42745 (< 1) |
| Gall vs No_gall | 114,625 | 4,827 | 2,039 | 1,403 | 48757 (< 2) |

Egg: *Diplolepis rosae* eggs removed from the female adults. Gland_Nov: salivary glands from the *D. rosae* larva collected in November. Gland_Oct: salivary glands from the *D. rosae* larva collected in October. Head: *D. rosae* female adult head. Larva_July: *D. rosae* larva removed from the gall in mid-July. Larva_Sept: *D. rosae* larva removed from the gall in early September. No_growth: sample set which does not correspond to the active gall growth stage (Egg + Head + Gland_Oct + Gland_Nov). Growth: sample set corresponding to the active gall growth stage (Larva_July + Larva_Sept + D_eglant, i.e. mid-July *D. eglanteriae* larva). No_gall: sample set which does not correspond to the whole gall stage (Egg + Head): Gall: sample set corresponding to the whole gall stage (Larva_July + Larva_Sept + Gland_Oct + Gland_Nov + D_eglant, mid-July *D. eglanteriae* larva). Total non-zero: number of genes that reads mapped at least one time. Up: number of up-regulated genes, i.e. whose expression is more in the first *D. rosae* tissue/stage compared to the second one in the first *D. rosae* tissue/stage compared to the second one. The number of up-regulated genes was defined by positive log2(read counts in the first tissue/read counts of the second tissue) (p adj < 0.05) (**Figures S11-S13**). Down: number of down-regulated genes, i.e. whose expression is less in the first *D. rosae* tissue/stage compared to the second one. The number of down-regulated genes was defined by negative log2(read counts in the first tissue/read counts of the second tissue) (p adj < 0.05) (**Figures S11-S13**). Outliers: number of genes where reads show outlier counts. Low counts (mean count <): number of genes where a number of mapped reads is less than the given threshold. The total number of examined genes was 125,626.

**Table S3.** Functional annotation and selection coefficients of the genes encoding proteins with leucine-rich repeats, plants cell wall degrading enzymes, and venom-like proteins that were overexpressed during gall formation from mid-July to early November in *Diplolepis rosae.*

| **Gene id** | **Annot lev** | **Func annot**  **eggNOG** | **MK**  **(*s*)** | **N**  **tr** | **L, aa** | **blastp sp align** | **blastp**  **match** | **% cov** | **% id** | **E value** | **tot sc** |
| --- | --- | --- | --- | --- | --- | --- | --- | --- | --- | --- | --- |
| g29097 | Hym | Leucine-  rich repeat | -3,34 (neg) | 2 | 776 | *Belonocnema* sp. | Slit homolog | 97 | 77.92 | 0 | 1068 |
| g39136  * | Hym | Leucine-  rich repeat | -1.27  (neg) | 2 | 498 | *Belonocnema* sp. | Slit homolog | 84 | 47.31 | 2e-142 | 446 |
| g51958  * | Hym | Leucine-  rich repeat | -1.19  (neg) | 1 | 723 | *Belonocnema* sp. | Podocan | 97 | 69.08 | 0 | 911 |
| g53074 | Hym | Leucine-  rich repeat | -0.80 (neg) | 2 | 1234 | *Belonocnema* sp. | Chaoptin-  Like | 100 | 85.33 | 0 | 2100 |
| g79908 | Hym | Leucine-  rich repeat | -0.92 (neg) | 2 | 247 | *Belonocnema* sp. | U2 small ribonucleo-  protein A’ | 100 | 90.69 | 3e-162 | 459 |
| g86458 | Hym | Leucine-  rich repeat | -1.29 (neg) | 2 | 298 | *Belonocnema* sp. | Peroxidasin-  like | 95 | 69.18 | 4e-129 | 380 |
| g86477 | Hym | Leucine-  rich repeat | -1.55 (neg) | 2 | 351 | *Belonocnema* sp. | Protein phosphatase 1 regulatory subunit 42-like | 100 | 72.75 | 0 | 533 |
| g87196 | Hym | Leucine-  rich repeat | -1.09 (neg) | 1 | 1351 | *Leptopilina* sp. | Chaoptin | 99 | 80.43 | 0 | 2149 |
| g108407* | Hym | Leucine-  rich repeat | -2.05 (neg) | 1 | 699 | *Leptopilina* sp. | Vasorin  -like | 34 | 66.39 | 2e-101 | 330 |
| g108407* | Hym | Leucine-  rich repeat | -2.05 (neg) | 1 | 699 | *Leptopilina* sp. | Neuronal protein 1-like | 34 | 66.39 | 6e-94 | 311 |
| g108411* | Hym | Leucine-  rich repeat | -1.20 (neg) | 1 | 133 | *Harpegnathos* sp. | Neuronal protein 2 | 79 | 67.92 | 9e-46 | 165 |
| g120908* | Hym | Leucine-  rich repeat | -1.69 (neg) | 2 | 568 | *Belonocnema* sp. | Carboxypeptidase N subunit 2-like | 95 | 65.81 | 0 | 701 |
| g120908* | Hym | Leucine-  rich repeat | -1.69 (neg) | 2 | 568 | *Nomia* sp. | Toll-like receptor 7 | 97 | 40.04 | 3e-125 | 389 |
| g122472 | Insect | Leucine-  rich repeat | -1.34 (neg) | 1 | 383 | *Lasius* sp. | Slit-like | 37 | 61.74 | 1e-50 | 186 |
| g122552 | Hym | Leucine-  rich repeat | -1.34 (neg) | 2 | 611 | *Leptopilina* sp. | Insulin-like growth factor-  binding protein | 100 | 78.57 | 0 | 993 |
| g122552 | Hym | Leucine-  rich repeat | -1.45 (neg) | 2 | 611 | *Belonocnema* sp. | Carboxy-  peptidase N subunit 2 | 100 | 84.90 | 0 | 1059 |
| g123069 | Hym | Leucine-  rich repeat | -0.75 (neg) | 2 | 902 | *Belonocnema* sp. | Fibronectin type-III domain containing protein | 99 | 89.64 | 0 | 1583 |
| g123069 | Hym | Leucine-  rich repeat | -0.75 (neg) | 2 | 902 | *Leptopilina* sp. | Vasorin | 98 | 86.49 | 0 | 1553 |
| g124026 | Hym | Leucine-  rich repeat | -0.74 (neg) | 1 | 927 | *Leptopilina* sp. | Follicle-  stimulating hormone receptor | 98 | 84.95 | 0 | 1612 |
| g31505 | GPA | Rhamno-  galacturonate lyase | -1.33 (neg) | 1 | 595 | *Belonocnema* sp. | Rhamno-  galacturonate lyase-like | 97 | 54.02 | 0 | 630 |
| g51222 | GPA | Pectate lyase | -1.17 (neg) | 1 | 337 | *Belonocnema* sp. | Pectin lyase-like | 89 | 48.68 | 6e-99 | 305 |
| g51223 | GPA | Pectate lyase | -1.22 (neg) | 1 | 264 | *Belonocnema* sp. | Pectin lyase-like | 99 | 50.00 | 4e-47 | 254 |
| g81883 | GPA | Cellulase | -1.43 (neg) | 3 | 373 | *Belonocnema* sp. | Endo  glucanase Z-like | 80 | 65.22 | 1e-139 | 410 |
| g87016 | GPA | Pectate lyase | -1.60 (neg) | 1 | 330 | *Belonocnema* sp. | Pectin lyase-like | 90 | 63.67 | 9e-134 | 393 |
| g94279 | GPA | Cellulase | -1.21 (neg) | 1 | 512 | *Belonocnema* sp. | Endo  glucanase Z-like | 60 | 71.61 | 2e-163 | 476 |
| g30759 | Hym | Venom protease-  like / Trypsin | -3.69 (neg) | 1 | 187 | *Leptopilina* sp. | Venom serine protease Bi-VSP-  like | 61 | 55.65 | 2e-32 | 130 |
| g30762 | Hym | Venom protease-  like / Trypsin | -1.84 (neg) | 1 | 245 | *Belonocnema* sp. | Venom serine protease Bi-VSP-  like | 40 | 72.28 | 2e-42 | 155 |
| g30762 | Hym | Venom protease-  like / Trypsin | -1.84 (neg) | 1 | 245 | *Leptopilina* sp. | Venom serine protease Bi-VSP-  like | 46 | 62.07 | 2e-40 | 153 |
| g49748 * | Hym | Venom acid phosphatase / Histidine phosphatase 2 | -1.26 (neg) | 1 | 302 | *Belonocnema* sp. | Venom acid phosphatase Acph-1-  like | 99 | 72.09 | 2e-162 | 467 |
| g49748  * | Hym | Venom acid phosphatase  / Histidine phosphatase 2 | -1.26 (neg) | 1 | 302 | *Leptopilina* sp. | Venom acid phosphatase Acph-1-  like | 99 | 67.00 | 5e-151 | 438 |
| g67338  * | Hym | Venom acid phosphatase  / Histidine phosphatase 2 | -1.43 (neg) | 1 | 375 | *Belonocnema* sp. | Venom acid phosphatase Acph-1-  like | 100 | 61.27 | 8e-170 | 489 |
| g67338  * | Hym | Venom acid phosphatase  / Histidine phosphatase 2 | -1.43 (neg) | 1 | 375 | *Leptopilina* sp. | Venom acid phosphatase Acph-1-  like | 99 | 56.23 | 3e-149 | 437 |
| g74434  * | Hym | Venom acid phosphatase  / Histidine phosphatase 2 | -1.44 (neg) | 1 | 192 | *Leptopilina* sp. | Venom acid phosphatase Acph-1-  like | 81 | 48.72 | 4e-45 | 163 |
| g74436  * | Hym | Venom acid phosphatase  / Histidine phosphatase 2 | -1.63 (neg) | 1 | 214 | *Belonocnema* sp. | Venom acid phosphatase Acph-1-  like | 57 | 56.10 | 9e-39 | 147 |
| g74436  * | Hym | Venom acid phosphatase  / Histidine phosphatase 2 | -1.63 (neg) | 1 | 214 | *Leptopilina* sp. | Venom acid phosphatase Acph-1-  like | 53 | 54.78 | 5e-37 | 147 |
| g60751 * | Hym | Tetraspanin | -0.78 (neut) | 2 | 290 | *Leptopilina* sp. | CD63 antigen-like | 100 | 82.07 | 3e-168 | 478 |
| g121032  * | Meta | Domain of unknown function | -1.85 (neg) | 1 | 119 | *Trichonephila clavipes* | Putative transposase | 85 | 57.84 | 5e-36 | 130 |
| g28790 | Hym | Phosphatidylethanolamine-binding protein | -1.40 (neg) | 3 | 208 | *Belonocnema*  sp. | Protein D2-like | 100 | 86.12 | 4e-134 | 385 |
| g53036 | Hym | Chitooligosaccharidolytic beta-N-acetylglucosaminidase | -3.36 (neg) | 2 | 599 | *Leptopilina* sp. | Chitooligosaccharidolytic beta-N-acetylglucosaminidase | 100 | 68.50 | 0 | 896 |
| g53036 | Hym | Chitooligosaccharidolytic beta-N-acetylglucosaminidase | -3.36 (neg) | 2 | 599 | *Nasonia vitripennis* | Chitooligosaccharidolytic beta-N-acetylglucosaminidase | 100 | 61.53 | 0 | 808 |
| g53036 | Hym | Chitooligosaccharidolytic beta-N-acetylglucosaminidase | -3.36 (neg) | 2 | 599 | *Venturia canescens* | Chitooligosaccharidolytic beta-N-acetylglucosaminidase | 96 | 62.76 | 0 | 806 |
| g90912 | Hym | Apyrase | -1.41 (neg) | 1 | 131 | *Belonocnema*  sp. | Soluble calcium-  activated nucleotidase 1 | 90 | 75.00 | 8e-58 | 194 |
| g90912 | Hym | Apyrase | -1.41 (neg) | 1 | 131 | *Leptopilina* sp. | Soluble calcium-  activated nucleotidase 1 | 90 | 67.50 | 1e-51 | 178 |
| g90912 | Hym | Apyrase | -1.41 (neg) | 1 | 131 | *Leptopilina* sp. | Apyrase | 90 | 69.17 | 4e-51 | 177 |
| g118636 | Hym | Inosine-  uridine preferring nucleoside hydrolase | -1.89 (neg) | 2 | 342 | *Belonocnema*  sp. | Probable uridine nucleosidase 2 | 97 | 63.06 | 2e-149 | 434 |
| g118639 | Hym | Lysozyme C | -1.56 (neg) | 1 | 144 | *Belonocnema*  sp. | Lysozyme C1-like | 96 | 74.82 | 7e-75 | 230 |
| g121541 * | Insect | Testicular haploid expressed repeat | -1.90 (neg) | 1 | 130 | *Leptopilina* sp. | Testicular haploid expressed protein-like | 79 | 69.90 | 5e-36 | 136 |
| g36882 | Hym | Group XII secretory phospho-  lipase A2 precursor | -0.80 (neg) | 1 | 264 | *Leptopilina* sp. | Group XII secretory phospho-  lipase A2 | 77 | 82.24 | 1e-126 | 369 |
| g36882 | Hym | Group XII secretory phospho-  lipase A2 precursor | -0.80 (neg) | 1 | 264 | *Belonocnema*  sp. | Group XII secretory phospho-  lipase A2 | 77 | 83.33 | 2e-124 | 363 |
| g36882 | Hym | Group XII secretory phospho-  lipase A2 precursor | -0.80 (neg) | 1 | 264 | *Nasonia vitripennis* | Group XII secretory phospho-  lipase A2 | 79 | 78.95 | 9e-121 | 353 |
| g95508 | Hym | Phospho-  lipase A2-like | -1.29 (neg) | 2 | 189 | *Belonocnema*  sp. | Phospho-  lipase A2-like | 95 | 62.43 | 7e-83 | 255 |
| g95508 | Hym | Phospho-  lipase A2-like | -1.29 (neg) | 2 | 189 | *Leptopilina* sp. | Phospho-  lipase A2-like | 97 | 59.46 | 4e-79 | 245 |
| g124077 * | Hym | Phospho-  lipase A2 | -1.23 (neg) | 1 | 230 | *Belonocnema*  sp. | Phospho-  lipase A2 | 86 | 76.50 | 4e-111 | 328 |
| g124077  * | Hym | Phospho-  lipase A2 | -1.23 (neg) | 1 | 230 | *Leptopilina* sp. | Phospho-  lipase A2-like | 84 | 73.71 | 6e-102 | 305 |
| g124077 * | Hym | Phospho-  lipase A2 | -1.23 (neg) | 1 | 230 | *Cyphomyrmex costatus* | Predicted: phospho-  lipase A2 A2-actitoxin-Usc2a | 83 | 64.62 | 9e-91 | 277 |
| g124077 * | Hym | Phospho-  lipase A2 | -1.23 (neg) | 1 | 230 | *Colletes gigas* | Phospho-  lipase A2 A2-actitoxin-Usc2a | 85 | 62.50 | 1e-90 | 276 |
| g53492  * | Hym | AB hydrolase superfamily: lipase family | -1.02 (neg) | 2 | 612 | *Leptopilina* sp. | Lipase-3-  like | 63 | 69.82 | 0 | 606 |
| g53493  * | Hym | AB hydrolase superfamily: lipase family | -0.68 (neut) | 2 | 429 | *Belonocnema*  sp. | Lipase-3-  like | 85 | 83.11 | 0 | 950 |
| g53493  * | Hym | AB hydrolase superfamily: lipase family | -0.68 (neut) | 2 | 429 | *Leptopilina* sp. | Lipase-3-  like | 99 | 71.96 | 0 | 659 |
| g71703 | Hym | AB hydrolase superfamily: lipase family | -1.46 (neg) | 2 | 448 | *Leptopilina* sp. | Lipase-3-  like | 93 | 67.38 | 0 | 613 |
| g71703 | Hym | AB hydrolase superfamily: lipase family | -1.46 (neg) | 2 | 448 | *Nasonia vitripennis* | Lipase-3-  like | 85 | 55.93 | 4e-151 | 447 |
| g71708  * | Hym | AB hydrolase superfamily: lipase family | -1.35 (neg) | 1 | 402 | *Belonocnema*  sp. | Lipase-3-  like | 96 | 54.64 | 1e-177 | 513 |
| g123552 | Hym | AB hydrolase superfamily: lipase family | -1.45 (neg) | 1 | 286 | *Leptopilina* sp. | Lipase-3-  like | 100 | 64.34 | 2e-139 | 410 |
| g52886 | Hym | Lipase | -1.40 (neg) | 1 | 433 | *Leptopilina* sp. | Pancreatic lipase-  related protein-1-  like | 86 | 59.36 | 2e-169 | 491 |
| g52886 | Hym | Lipase | -1.40 (neg) | 1 | 433 | *Orussus abietinus* | Pancreatic triacyl  glycerol lipase | 84 | 54.59 | 6e-150 | 441 |
| g74954  * | Hym | Lipase | -1.07 (neg) | 3 | 313 | *Belonocnema*  sp. | Lipase member H | 100 | 92.97 | 0 | 617 |
| g74954  * | Hym | Lipase | -1.07 (neg) | 3 | 313 | *Leptopilina* sp. | Phospho-  lipase A1 memeber A | 100 | 88.33 | 0 | 588 |
| g74954  * | Hym | Lipase | -1.07 (neg) | 3 | 313 | *Odontomachus brunneus* | Pancreatic lipase-  related protein-2-  like | 100 | 84.98 | 0 | 573 |
| g94397  * | Hym | Lipase | -1.46 (neg) | 1 | 355 | *Belonocnema*  sp. | Pancreatic triacyl  glycerol lipase | 93 | 67.98 | 4e-175 | 500 |
| g94397  * | Hym | Lipase | -1.90 (neg) | 1 | 355 | *Leptopilina* sp. | Pancreatic lipase-  related protein-1-  like | 89 | 45.99 | 4e-93 | 293 |
| g120373 | Hym | Lipase | -1.46 (neg) | 1 | 388 | *Belonocnema*  sp. | Pancreatic triacyl  glycerol lipase-like | 96 | 78.46 | 0 | 630 |
| g120373 | Hym | Lipase | -1.46 (neg) | 1 | 388 | *Leptopilina* sp. | Pancreatic triacyl  glycerol lipase-like | 98 | 72.92 | 0 | 600 |
| g124088 | Hym | Animal haem peroxidase | -1.39 (neg) | 1 | 698 | *Leptopilina* sp. | Peroxidase-  like | 100 | 68.62 | 0 | 998 |
| g85953 | Hym | Animal haem peroxidase | -1.61 (neg) | 1 | 1312 | *Belonocnema*  sp. | Peroxidase | 51 | 89.36 | 0 | 1321 |
| g118333 | Hym | Animal haem peroxidase | -1.22 (neg) | 1 | 693 | *Leptopilina* sp. | Peroxidasin homolog | 99 | 83.21 | 0 | 1238 |
| g119756  * | Hym | Animal haem peroxidase | -1.33 (neg) | 1 | 1280 | *Belonocnema*  sp. | Peroxidasin | 100 | 81.74 | 0 | 2259 |

Annot lev: annotation level. Hym: Hymenoptera. GPA: Gammaproteobacteria. Insect: Insecta. Met: Metazoa. Func annot eggNOG: functional annotation provided by eggnog-mapper v. 2.1.7 (Cantalapiedra et al., 2021; Huerta-Cepas et al., 2021). MK (*s*): selection coefficient estimated by SnIPRE, McDonald-Kreitman type analysis (Eilertson et al., 2012); neg: negative selection; neut: neutral selection. N tr: number of expressed gene transcripts. L, aa: protein length, amino acids. Blastp sp align: taxonomic name of the species showing protein sequences matching to the query *Diplolepis rosae* sequence. % cov: query cover, proportion of the query *D. rosae* sequence that is aligned with the database sequence. % id: percent of identity between the query *D. rosae* sequence and the database sequence. E val: expectation value of the alignment between the query *D. rosae* sequence and the database sequence. Tot sc: total score reflecting the strength of the match between the query *D. rosae* sequence and the database sequence. The query *D. rosae* protein and the database protein showing E value < 10e-06 and total score < 50 were considered homologous (Pearson, 2013). Blue columns indicate genes encoding proteins with leucine rich repeat (Zhao et al., 2015). Yellow columns indicate genes encoding plant cell wall degrading enzymes (Hearn et al. 2019). Red columns indicate genes encoding proteins found in the *D. rosae* venom glands (Cambier et al., 2019). Green columns indicate the *D. rosae* genes orthologous to the genes overexpressed in the *B. pallida* venom glands (Cambier et al., 2019). The asterisks (*) denotes genes highly expressed in the mid-July and early September *D. rosae* larvae.

**Table S4.** Characteristics of *Diplolepis rosae* and *Diplolepis eglanteriae* samples used in Illumina sequencing.

| Species | Sample name | Stage | Number of individuals per sample | Tissue | Sex | Host plant | Isolation source | Collection date | Latitude | Longitude |
| --- | --- | --- | --- | --- | --- | --- | --- | --- | --- | --- |
| *Diplolepis rosae* | ESE-078 | Imago | 6 | Whole individual | Female | *Rosa canina* | Gall | 2020-05-12 | 48.70199 | 2.173458 |
| *Diplolepis rosae* | ESE-082 | Imago | 7 | Whole individual | Female | *Rosa canina* | Gall | 2020-05-13 | 48.70371 | 2.083596 |
| *Diplolepis rosae* | ESE-117 | Imago | 8 | Whole individual | Female | *Rosa canina* | Gall | 2020-04-04 | 45.75876 | 6.159383 |
| *Diplolepis rosae* | ESE-126 | Imago | 8 | Whole individual | Female | *Rosa canina* | Gall | 2020-06-14 | 47.24930 | 6.073626 |
| *Diplolepis rosae* | ESE-219 | Imago | 30 | Whole individual | Female | *Rosa canina* | Gall | 2020-08-10 | 46.42217 | 1.353230 |
| *Diplolepis rosae* | ESE-288 | Larva | 7 | Whole individual | Female | *Rosa canina* | Gall | 2020-08-26 | 47.32478 | 3.792692 |
| *Diplolepis rosae* | ESE-312 | Larva | 10 | Whole individual | Female | *Rosa canina* | Gall | 2020-09-02 | 50.00647 | 4.743896 |
| *Diplolepis rosae* | ESE-330 | Larva | 20 | Whole individual | Female | *Rosa canina* | Gall | 2020-09-05 | 44.52756 | 5.016034 |
| *Diplolepis rosae* | ESE-349 | Larva | 10 | Whole individual | Female | *Rosa canina* | Gall | 2020-09-08 | 45.62013 | 3.057751 |
| *Diplolepis rosae* | ESE-464 | Larva | 10 | Whole individual | Female | *Rosa canina* | Gall | 2020-09-23 | 43.44791 | 6.239238 |
| *Diplolepis rosae* | ESE-559 | Larva | 12 | Whole individual | Female | *Rosa canina* | Gall | 2020-09-10 | 43.96794 | 3.405437 |
| *Diplolepis rosae* | ESE-576 | Larva | 10 | Whole individual | Female | *Rosa canina* | Gall | 2020-10-12 | 43.03309 | 1.003795 |
| *Diplolepis rosae* | ESE-580 | Larva | 10 | Whole individual | Female | *Rosa canina* | Gall | 2020-10-09 | 42.70232 | 2.564203 |
| *Diplolepis rosae* | ESE-608 | Larva | 8 | Whole individual | Female | *Rosa canina* | Gall | 2020-10-20 | 44.58980 | -0.454849 |
| *Diplolepis rosae* | ESE-623 | Larva | 7 | Whole individual | Female | *Rosa canina* | Gall | 2020-10-20 | 43.65517 | -1.180453 |
| *Diplolepis rosae* | ESE-630 | Larva | 10 | Whole individual | Female | *Rosa canina* | Gall | 2020-10-20 | 45.15035 | 0.529976 |
| *Diplolepis rosae* | ESE-652 | Larva | 5 | Whole individual | Female | *Rosa canina* | Gall | 2020-12-27 | 47.30910 | -3.067183 |
| *Diplolepis eglanteriae* | ESE-54 | - | 1 | Whole individual | Female | *Rosa sp.* | Gall | 2019-10-  09 | 48.421111 | 0.923611 |
| *Diplolepis eglanteriae* | ESE-281 | Imago | 1 | Whole individual | Female | *Rosa canina* | Gall | 2020-08-  27 | 47.286478 | 4.7273 |

**Table S5.** Quality control of *Diplolepis rosae* RNAseq libraries used in RNAseq analysis.

| **Lib** | **Num pairs, Mln** | **Num bases, Gb** | **Effect, %** | **Error rate, %** | **Q20, %** | **Q30, %** | **GC cont, %** | **Per base GC cont** | **Per base seq cont** | **Per base**  **N cont** | **Seq dup levels** | **Over**  **seq** | **Adapt** |
| --- | --- | --- | --- | --- | --- | --- | --- | --- | --- | --- | --- | --- | --- |
| Head_1 | 106.0 | 15.9 | 98.82 | 0.03 | 97.69 | 93.43 | 34.19 | Fail | Fail | Pass | Fail | Warn | Pass |
| Head_2 | 106.0 | 15.9 | 98.33 | 0.03 | 97.52 | 93.00 | 34.26 | Fail | Fail | Pass | Fail | Warn | Pass |
| Egg_1 | 99.66 | 14.9 | 97.64 | 0.03 | 97.34 | 92.79 | 35.52 | Fail | Fail | Pass | Fail | Warn | Pass |
| Egg_3 | 91.81 | 13.8 | 98.21 | 0.03 | 97.73 | 93.47 | 35.37 | Pass | Fail | Pass | Fail | Warn | Pass |
| Egg_3 | 11.74 | 1.76 | 98.12 | 0.03 | 96.59 | 90.82 | 35.15 | Pass | Fail | Pass | Fail | Warn | Pass |
| Egg_4 | 97.91 | 14.7 | 98.76 | 0.03 | 97.45 | 93.00 | 35.99 | Pass | Fail | Pass | Fail | Warn | Pass |
| Larva_  July_1 | 7.72 | 11.6 | 98.76 | 0.03 | 96.17 | 90.06 | 40.74 | Fail | Fail | Pass | Fail | Warn | Pass |
| Larva_  July_1 | 94.44 | 14.2 | 98.75 | 0.03 | 97.45 | 92.93 | 40.96 | Fail | Fail | Pass | Fail | Warn | Pass |
| Larva_  July_2 | 99.76 | 15.0 | 98.88 | 0.03 | 97.78 | 93.71 | 40.42 | Warn | Fail | Pass | Fail | Warn | Pass |
| Larva_  July_3 | 81.15 | 12.2 | 98.92 | 0.03 | 97.65 | 93.40 | 39.06 | Pass | Fail | Pass | Fail | Warn | Pass |
| Larva_  July_3 | 24.50 | 3.68 | 98.93 | 0.03 | 96.47 | 90.69 | 38.86 | Pass | Fail | Pass | Fail | Warn | Pass |
| Larva_  July_4 | 28.50 | 4.28 | 98.79 | 0.03 | 97.37 | 92.89 | 42.47 | Fail | Fail | Pass | Fail | Warn | Pass |
| Larva_  July_4 | 79.72 | 12.0 | 98.81 | 0.03 | 97.79 | 93.72 | 42.56 | Fail | Fail | Pass | Fail | Warn | Pass |
| Larva_Sept_1 | 18.61 | 2.79 | 98.92 | 0.03 | 97.47 | 93.05 | 36.95 | Fail | Fail | Pass | Fail | Warn | Pass |
| Larva_Sept_1 | 89.26 | 13.4 | 98.90 | 0.03 | 97.59 | 93.24 | 37.00 | Fail | Fail | Pass | Fail | Warn | Pass |
| Larva_Sept_2 | 98.36 | 14.8 | 98.82 | 0.03 | 97.37 | 92.80 | 36.77 | Fail | Fail | Pass | Fail | Warn | Pass |
| Larva_Sept_3 | 103.8 | 15.6 | 98.61 | 0.03 | 97.68 | 93.36 | 37.31 | Warn | Fail | Pass | Fail | Warn | Pass |
| Larva_Sept_4 | 16.66 | 2.50 | 98.74 | 0.03 | 97.44 | 93.01 | 37.67 | Warn | Fail | Pass | Fail | Warn | Pass |
| Larva_Sept_4 | 91.55 | 13.7 | 98.72 | 0.03 | 97.57 | 93.20 | 37.72 | Warn | Fail | Pass | Fail | Warn | Pass |
| Gland_Oct_1 | 6.79 | 10.2 | 99.00 | 0.03 | 96.55 | 90.81 | 36.89 | Warn | Fail | Pass | Fail | Warn | Pass |
| Gland_Oct_1 | 96.56 | 14.5 | 99.00 | 0.03 | 97.70 | 93.44 | 37.14 | Warn | Fail | Pass | Fail | Warn | Pass |
| Gland_Oct_2 | 81.95 | 12.3 | 98.54 | 0.03 | 97.58 | 93.25 | 35.34 | Fail | Fail | Pass | Fail | Warn | Pass |
| Gland_Oct_2 | 24.36 | 3.65 | 98.52 | 0.03 | 96.57 | 90.90 | 35.05 | Fail | Fail | Pass | Fail | Warn | Pass |
| Gland_Oct_3 | 115.18 | 17.3 | 99.27 | 0.03 | 97.82 | 93.68 | 39.18 | Warn | Fail | Pass | Fail | Warn | Pass |
| Gland_Oct_4 | 82.06 | 12.3 | 98.71 | 0.03 | 96.63 | 91.13 | 38.17 | Warn | Fail | Pass | Fail | Warn | Pass |
| Gland_Oct_4 | 28.17 | 4.23 | 98.80 | 0.03 | 96.96 | 91.79 | 38.30 | Warn | Fail | Pass | Fail | Warn | Pass |
| Gland_Nov_1 | 122.9 | 18.4 | 99.29 | 0.03 | 97.64 | 93.25 | 38.11 | Warn | Fail | Pass | Fail | Pass | Pass |
| Gland_Nov_2 | 134.7 | 20.2 | 99.18 | 0.03 | 97.62 | 93.20 | 37.96 | Pass | Fail | Pass | Fail | Pass | Pass |
| Gland_Nov_3 | 64.96 | 9.74 | 98.67 | 0.03 | 97.54 | 93.13 | 38.49 | Pass | Fail | Pass | Fail | Pass | Pass |
| Gland_Nov_3 | 82.02 | 12.3 | 98.74 | 0.03 | 96.85 | 91.68 | 38.43 | Pass | Fail | Pass | Fail | Pass | Pass |
| Gland_Nov_4 | 83.21 | 12.5 | 99.02 | 0.03 | 97.63 | 93.34 | 38.20 | Warn | Fail | Pass | Fail | Pass | Pass |
| Gland_Nov_4 | 29.86 | 4.48 | 99.06 | 0.03 | 96.38 | 90.50 | 37.99 | Warn | Fail | Pass | Fail | Pass | Pass |
| D_eglant_2 | 83.89 | 10.6 | 98.43 | 0.03 | 97.07 | 92.12 | 38.22 | Warn | Fail | Pass | Fail | Warn | Pass |
| D_eglant_2 | 67.21 | 10.1 | 98.32 | 0.03 | 97.68 | 93.42 | 38.27 | Warn | Fail | Pass | Fail | Warn | Pass |
| D_eglant_4 | 60.33 | 9.05 | 99.07 | 0.03 | 97.77 | 93.67 | 37.13 | Warn | Fail | Pass | Fail | Warn | Pass |
| D_eglant_4 | 97.28 | 14.6 | 99.10 | 0.03 | 97.14 | 92.31 | 37.08 | Warn | Fail | Pass | Fail | Warn | Pass |

Head: *Diplolepis rosae* female adult head. Egg: *D. rosae* eggs removed from the female adults. Larva_July: *D. rosae* early larva removed from the gall in mid-July. Larva_Sept: *D. rosae* larva removed from the gall in early September. Gland_Oct: salivary glands from the *D. rosae* larva collected in October. Gland_Nov: salivary glands from the *D. rosae* larva collected in November. D_eglant: *D. eglanteriae*. Lib: RNAseq library. Num pairs: number of pairs of 150-bp pair-end reads. Num bases: total number of bases in the library. Effect: effective rate (100%*(clean reads / raw reads)). Error rate: base error rate. Q20: 100%*(base count of Qphred > 20 / total base count). Q30: 100%*(base count of Qphred > 30 / total base count). Q20 shows an error probability < 1/100. Q30 shows an error probability < 1/1000. GC cont: 100%*((G and C base count) / total base count). Per base GC cont: presence of contaminant sequences (Pass: GC distribution is near to theoretical normal distribution of an organism; Fail: presence of contaminant or overexpressed sequences). Per base seq cont: per base sequence content (always Fail in the RNA sequencing because of using random hexamer primers). Per base N cont: number of unidentified bases (Pass: the number of Ns is negligible). Seq dup levels: presence of duplicated sequences (Pass: < 20%; Warn (Warning): > 20%; Fail: > 50%) being overrepresented in the case of too little initial RNA quantity or too many PCR cycles. Over seq: sequences (at least 20 bp) occurring in more than 0.1% of the total number of sequences (overexpressed gene or adapter). Adapt: adapter sequences (Pass: removed by Novogene).

**Figures**

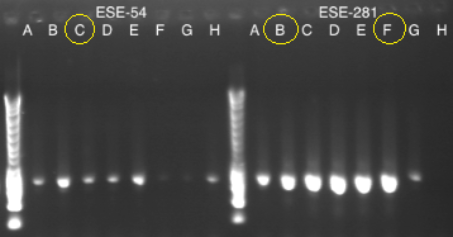

**Figure S1. Verification of PCR product (gene encoding cytochrome oxidase subunit I, COI) for further identification of *Diplolepis eglanteriae*.** ESE-54-C, ESE-281-B, and ESE-281-F were used for COI sequencing. ESE-54 and ESE-281-B were used for further Illumina sequencing.

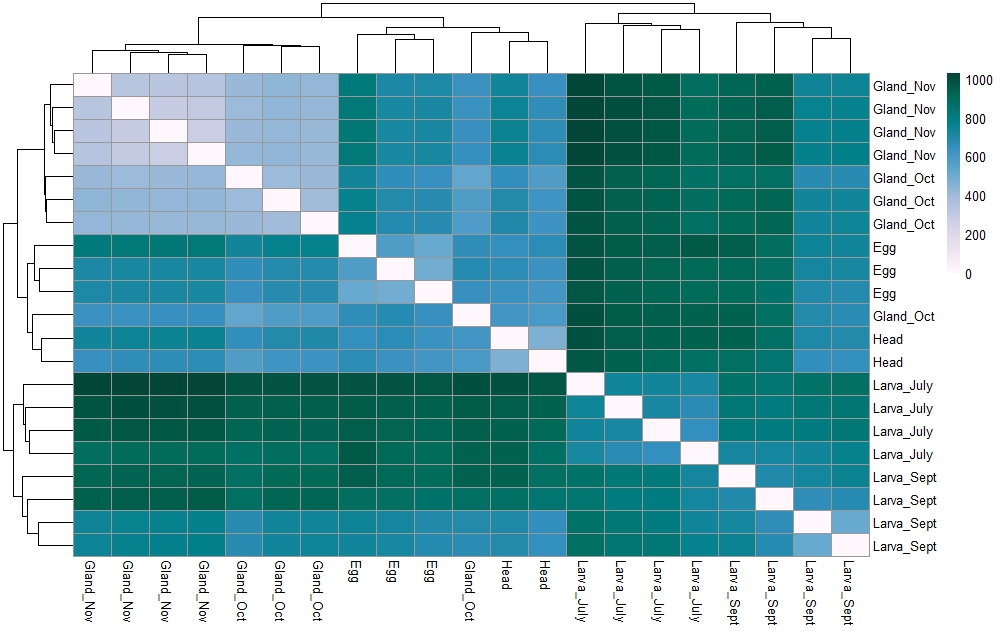

**Figure S2**. **Pairwise Euclidean distances between the *Diplolepis rosae* samples used in the relative differential gene expression analysis.** Egg: egg removed from a female adult. Larva_July: mid-July larva. Larva_Sept: early September larva. Gland_Oct: October larva salivary glands. Gland_Nov: November larva salivary glands. Head: female adult head. The clustering is based on the distances between the rows/columns of the distance matrix.

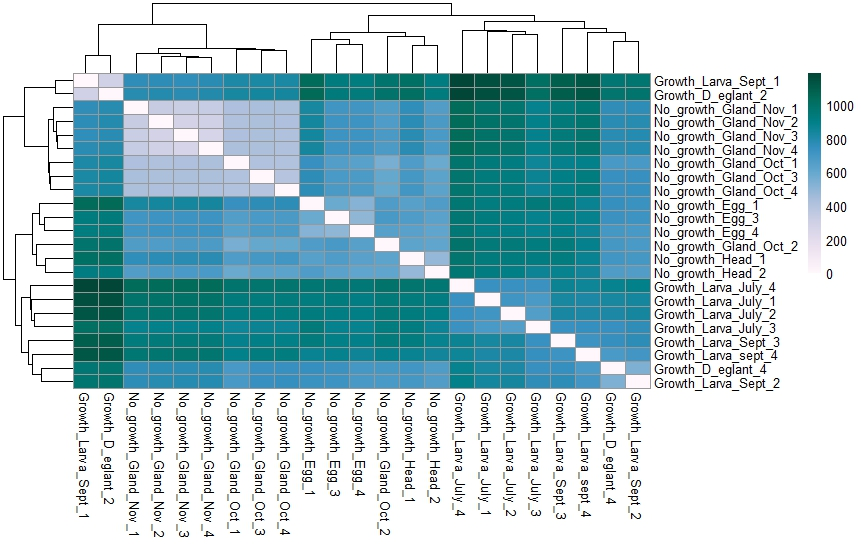

**Figure S3**. **Pairwise Euclidean distances between the No_growth’ and ‘Growth’ *Diplolepis rosae* samples used in the relative differential gene expression analysis.** No_growth: sample set that does not correspond to the active gall growth stage (egg removed from a female adult + female adult head + October larva salivary gland + November larva salivary gland). Growth: sample set corresponding to the active gall growth stage (mid-July larva + early September larva + mid-July *Diplolepis eglanteriae* larva). The clustering is based on the distances between the rows/columns of the distance matrix.

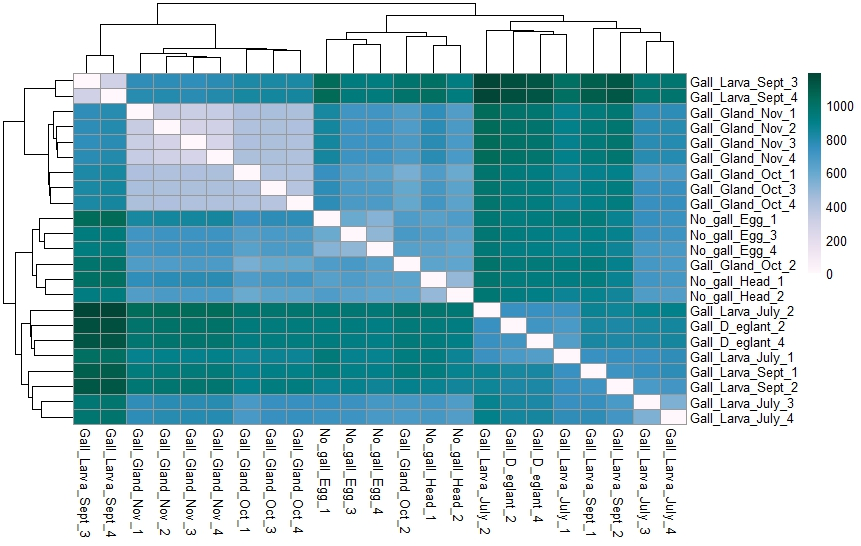

**Figure S4**. **Pairwise Euclidean distances between the No_gall’ and ‘Gall’ *Diplolepis rosae* samples used in the relative differential gene expression analysis**. No_gall: sample set which does not correspond to the whole gall stage (egg removed from a female adult + female adult head): Gall: sample set corresponding to the whole gall stage (mid-July larva + early September larva + mid-July *D. eglanteriae* larva + October salivary gland + November salivary gland). The clustering is based on the distances between the rows/columns of the distance matrix.

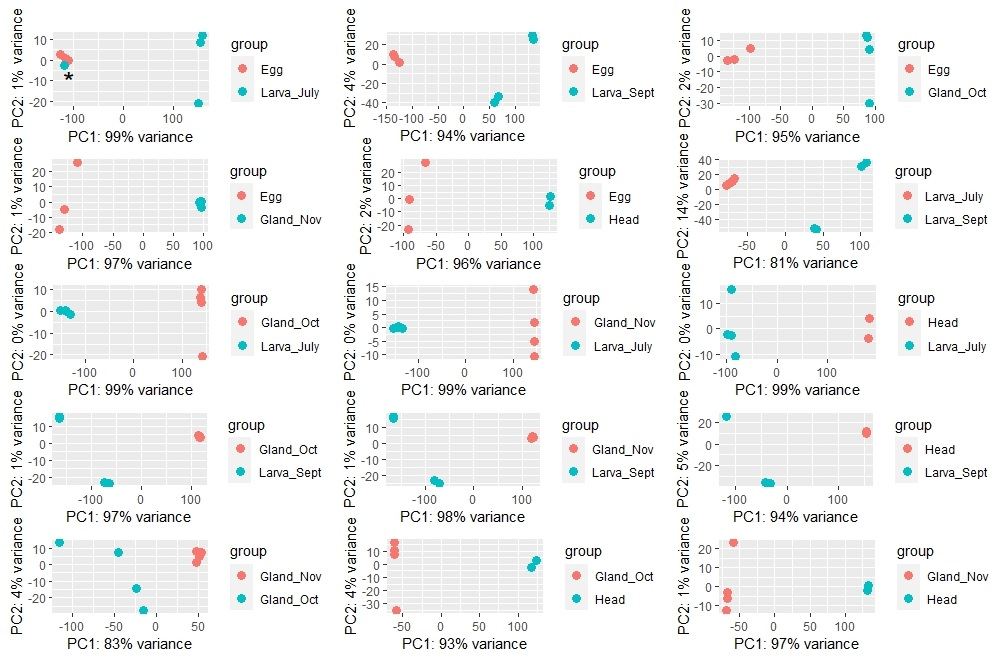

**Figure S5.** **Percentage of dispersion explained by the first (PC1) and the second (PC2) components after performing principal component analysis of the *Diplolepis rosae* samples used in the relative differential gene expression analysis**. Egg: egg removed from a female adult. Larva_July: mid-July larva. Larva_Sept: early September larva. Gland_Oct: October larva salivary glands. Gland_Nov: November larva salivary glands. Head: female adult head. *: sample removed from the analysis.

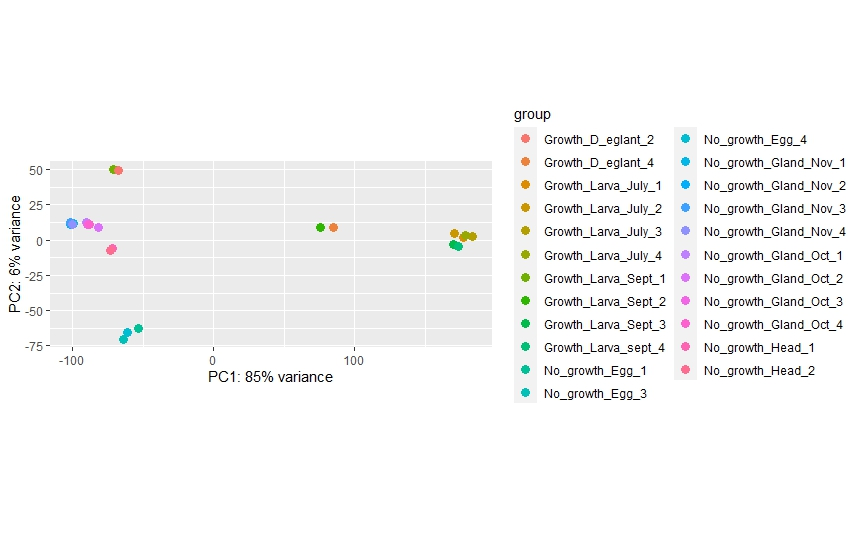

**Figure S6.** **Percentage of dispersion explained by the first (PC1) and the second (PC2) components after performing principal component analysis of the ‘No_growth’ and ‘Growth’ *D. rosae* samples used in the relative differential gene expression analysis**. No_growth: sample set that does not correspond to the active gall growth stage (egg removed from a female adult + female adult head + October larva salivary gland + November larva salivary gland). Growth: sample set corresponding to the active gall growth stage (mid-July larva + early September larva + mid-July *Diplolepis eglanteriae* larva).

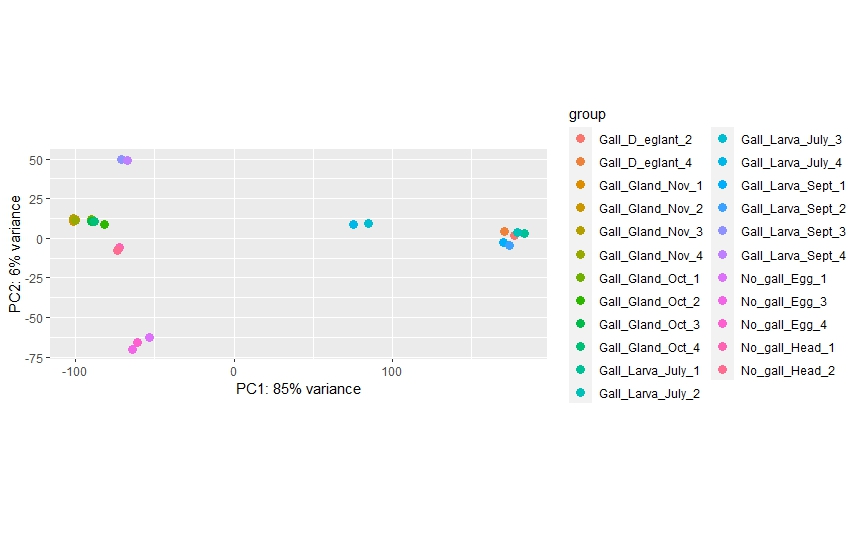

**Figure S7. Percentage of dispersion explained by the first (PC1) and the second (PC2) components after performing principal component analysis of the ‘No_gall’ and ‘Gall’ *Diplolepis rosae* samples used in the relative differential gene expression analysis**. No_gall: sample set which does not correspond to the whole gall stage (egg removed from a female adult + female adult head): Gall: sample set corresponding to the whole gall stage (mid-July larva + early September larva + mid-July *Diplolepis eglanteriae* larva + October salivary gland + November salivary gland).

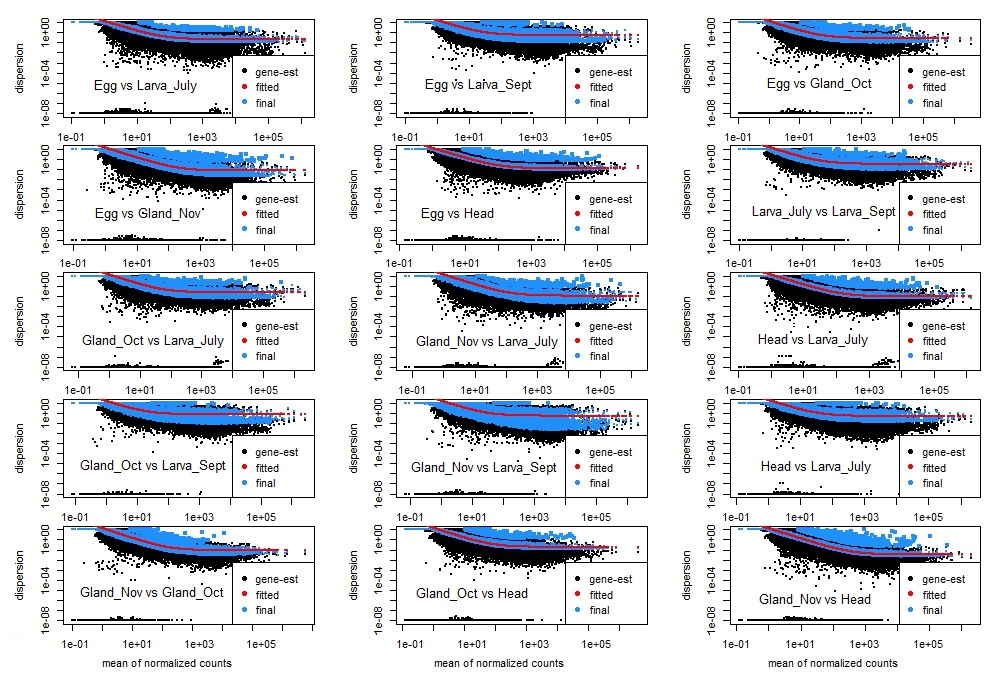

**Figure S8**. **Dispersion estimates provided by the pairwise relative differential gene expression analysis of the *Diplolepis rosae* samples**. gene-est (black points): gene estimation; dispersion values calculated for normalized mean counts of reads mapped to each gene. fitted (red line): fitting curve constructed according to a generalized linear model used in the analysis. final (blue points): shrank gene-wise dispersion estimates towards values predicted by the fitting curve. Egg: egg removed from a female adult. Larva_July: mid-July larva. Larva_Sept: early September larva. Gland_Oct: October larva salivary glands. Gland_Nov: November larva salivary glands. Head: female adult head.

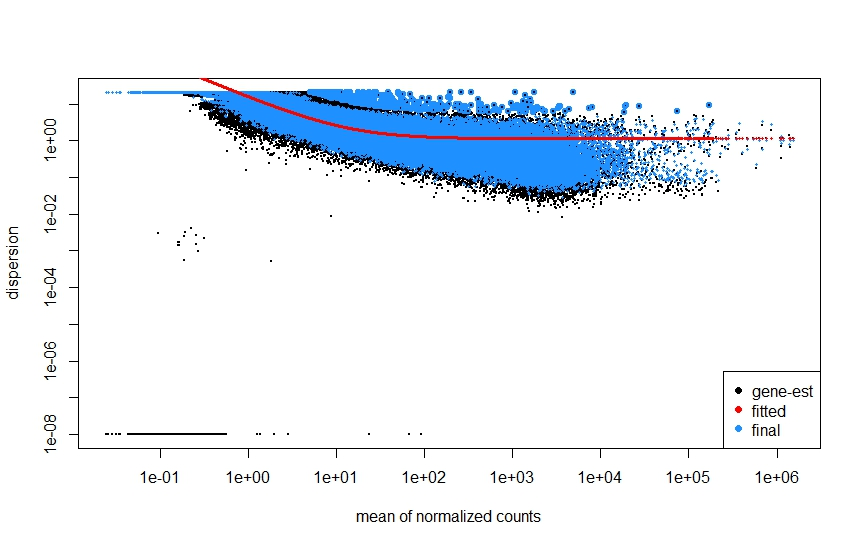

**Figure S9**. **Dispersion estimates provided by the relative differential gene expression analysis of the sample pair ‘No_growth *Diplolepis rosae* - Growth *Diplolepis rosae*’**. No_growth: sample set that does not correspond to the active gall growth stage (egg removed from a female adult + female adult head + October larva salivary gland + November larva salivary gland). Growth: sample set corresponding to the active gall growth stage (mid-July larva + early September larva + mid-July *Diplolepis eglanteriae* larva). gene-est (black points): gene estimation; dispersion values calculated for normalized mean counts of reads mapped to each gene. fitted (red line): fitting curve constructed according to a generalized linear model used in the analysis. final (blue points): shrank gene-wise dispersion estimates towards values predicted by the fitting curve.

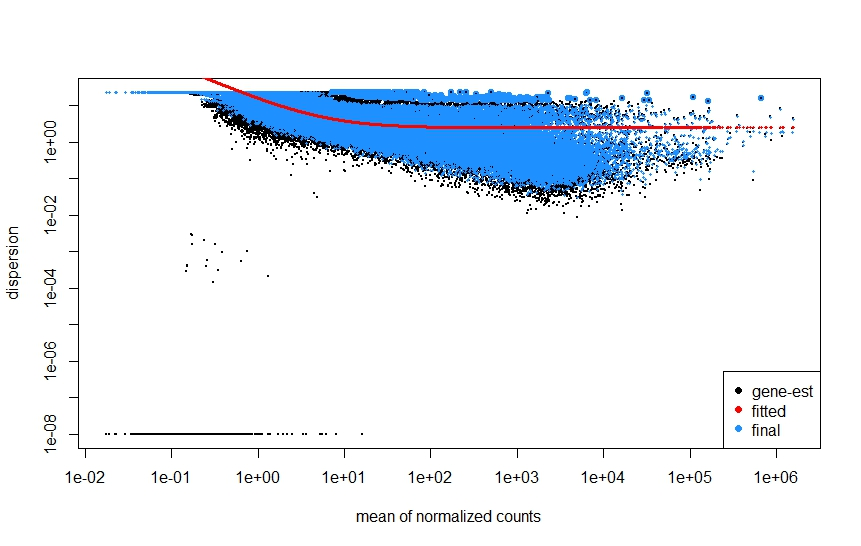

**Figure S10. Dispersion estimates provided by the relative differential gene expression analysis of the sample pair ‘No_gall *Diplolepis rosae* - Gall *Diplolepis. rosae*’**. No_gall: sample set which does not correspond to the whole gall stage (egg removed from a female adult + female adult head): Gall: sample set corresponding to the whole gall stage (mid-July larva + early September larva + mid-July *Diplolepis eglanteriae* larva + October salivary gland + November salivary gland). gene-est (black points): gene estimation; dispersion values calculated for normalized mean counts of reads mapped to each gene. fitted (red line): fitting curve constructed according to a generalized linear model used in the analysis. final (blue points): shrank gene-wise dispersion estimates towards values predicted by the fitting curve.

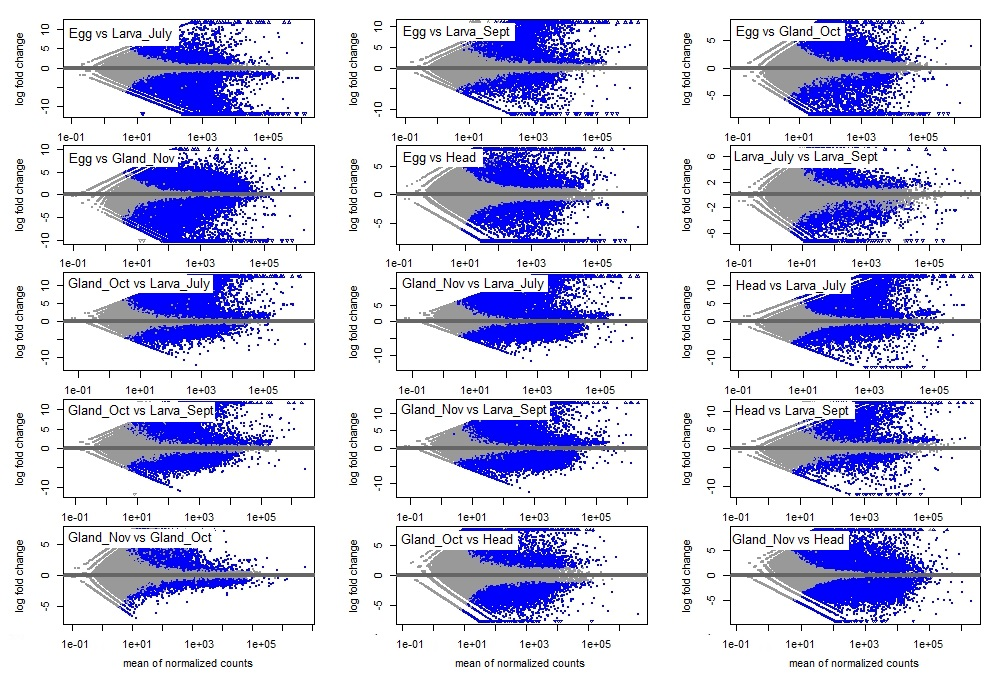

**Figure S11**. **Log2 fold changes of the gene expression levels calculated for each gene over the mean of normalized counts for all the samples used in the pairwise relative differential expression analysis**. Blue points show the values with the adjusted p-value < 0.1. Log2 fold change is calculated as log2 (normalized mean counts of reads mapped to a gene of sample 1 / normalized mean counts of reads mapped to a gene of sample 2). Egg: egg removed from a female adult. Larva_July: mid-July larva. Larva_Sept: early September larva. Gland_Oct: October larva salivary glands. Gland_Nov: November larva salivary glands. Head: female adult head.

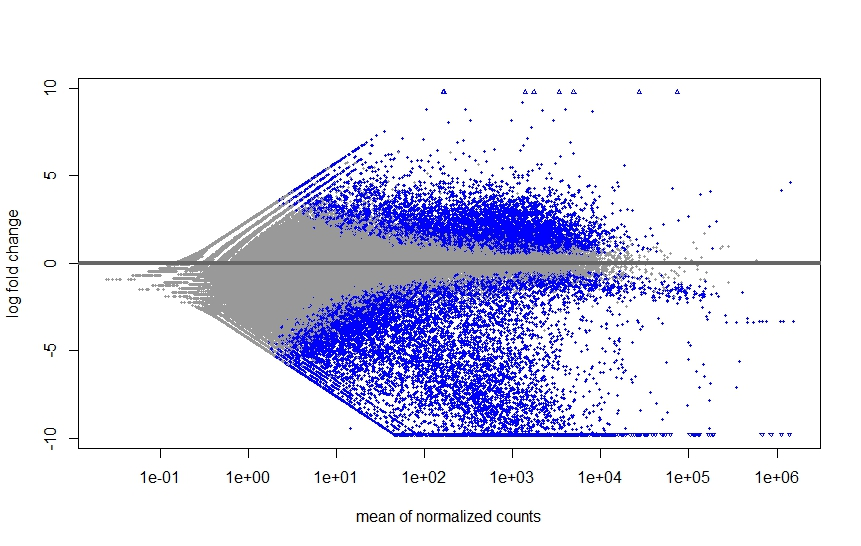

**Figure S12**. **Log2 fold changes of the gene expression levels calculated for each gene over the mean of normalized counts for all the ‘No_growth’ and ‘Growth’ *Diplolepis rosae* samples used in the relative differential expression analysis**. Blue points show the values with the adjusted p-value < 0.1. Log2 fold change is calculated as log2 (normalized mean counts of reads mapped to a gene of the ‘No_growth’ sample set / normalized mean counts of reads mapped to a gene of the ‘Growth’ sample set). No_growth: sample set which does not correspond to the active gall growth stage (egg removed from a female adult + female adult head + October larva salivary gland + November larva salivary gland). Growth: sample set corresponding to the active gall growth stage (mid-July larva + early September larva + mid-July *Diplolepis eglanteriae* larva).

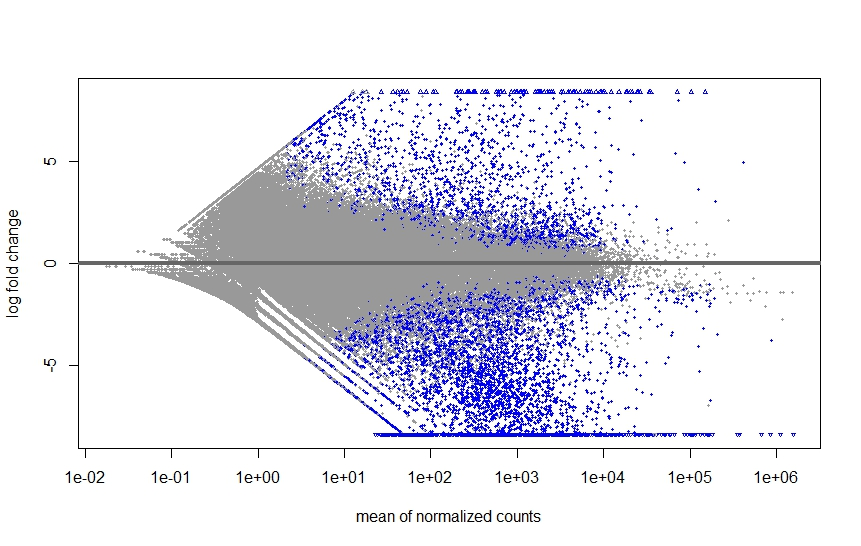

**Figure S13**. **Log2 fold changes of the gene expression levels calculated for each gene over the mean of normalized counts for all the ‘No_gall’ and ‘Gall’ *Diplolepis rosae* samples used in the relative differential expression analysis**. Blue points show the values with the adjusted p-value < 0.1. Log2 fold change is calculated as log2 (normalized mean counts of reads mapped to a gene of the ‘No_gall’ sample set / normalized mean counts of reads mapped to a gene of the ‘Gall’ sample set). No_gall: sample set which does not correspond to the whole gall stage (egg removed from a female adult + female adult head): Gall: sample set corresponding to the whole gall stage (mid-July larva + early September larva + mid-July *Diplolepis eglanteriae* larva + October salivary gland + November salivary gland).

**CODES**

**Code S1.** Variant calling

bowtie2-build reference_genome.fasta reference_genome

bowtie2 -x reference_genome -1 clean_reads_1.fastq.gz -2 clean_reads_2.fastq.gz -S alignment.sam

samtools view -S -b alignment.sam > alignment.bam

samtools sort alignment.bam -o alignment_sorted.bam

samtools stats alignment_sorted.bam ⇒ make after alignment

samtools depth  alignment_sorted.bam  |  awk '{sum+=$3} END { print "Average = ",sum/NR}' ⇒ make after alignment

samtools index -b alignment_sorted.bam

gatk AddOrReplaceReadGroups -I alignment_sorted.bam -O alignment_sorted_rg.bam -LB lib -PL ILLUMINA -PU unit -SM sample_name

samtools index -b alignment_sorted_rg.bam

samtools faidx reference_genome.fasta

samtools dict reference_genome.fasta

gatk HaplotypeCaller -I alignment_sorted_rg.bam -O genome_variant.gvcf -R reference_genome.fasta --emit-ref-confidence GVCF

gatk CombineGVCFs -R reference_genome.fasta -V genome_variant_1.gvcf -V genome_variant_2.gvcf … -V genome_variant_n.gvcf -O combined_genome_variant.gvcf

gatk GenotypeGVCFs -R reference_genome.fasta -V combined_genome_variant.gvcf  -O variant_call_file.vcf

**Code S2.** Identification of genes under selection (McDonald–Kreitman test)

java -Xmx100g -jar [beagle.22Jul22.46e.jar](http://faculty.washington.edu/browning/beagle/beagle.22Jul22.46e.jar) gt=variant_call_file.vcf out=phased_variant_call_file ne=500 window=50

bcftools filter -i 'TYPE="snp"' variant_call_file.vcf -Ov > vcf_snp.vcf

bcftools view --max-alleles 2 --exclude-types indels vcf_snp.vcf -Ov > vcf_snp_2all.vcf

samtools faidx reference.fasta

bgzip phased_variant_call_file.vcf

tabix phased_variant_call_file.vcf.gz

vcf2fasta.py -f reference.fasta -v phased_variant_call_file.vcf.gz -g annotaed_masked_from_transposons_file.gff3 -b -e CDS

MKtest -i alignment.fasta -I 1 1 (infoseq alignment.fasta | tail -1 | awk '{print $6}') -n N

*-n: number of haplotypes (N = 34 in D. rosae)*

polydNdS -i alignment.fasta -I 1 1 (infoseq $file | tail -1 | awk '{print $6}') -O N

*-O: outgroup individual number (N = 35 corresponding to Diplolepis eglanteriae)*

*Data to SnIPRE (Eilertson et al. 2012) script (R):*

  geneID FR PR FS PS     Trepl    Tsil nout npop

1  g1000  0  0  0  0  409.167  91.8333   10   34

2  g1001  0  0  0  0  163.500  40.5000   10   34

3  g1002  0  7  0  3 1365.280 332.7220   10   34

4  g1003  0  2  0  2 1295.720 294.2780   10   34

5  g1004  0  4  0  1  820.648 193.3520   10   34

6  g1005  0  2  0  1 2015.020 477.9810   10   34

…

```
## Part (1)  Empirical Bayes Implementation  (lme4 package, SnIPRE_source.R)
## Part (2)  Bayesian Implementation (R2WinBUGS package, B_SnIPRE_source.R, and WinBUGS or OpenBUGS) necessary
setwd("~/Dropbox/SnIPRE_code_JAGS")

#################################################################
## Part (1)  Empirical Bayes Implementation  (lme4 package)
#################################################################

source("SnIPRE_source.R")
source("my.jags2.R")
library(lme4)
library(R2jags)
library(arm)

data <- read.table("SnIPRE_table.txt", header = TRUE)  # sample data set

#SnIPRE <-function(mydata)
# mydata: name of data set;
# mydata must have a header with the following columns: PS, PR, FS, FR, npop, nout, Tsil, Trepl (no particular order)
# outputs 2 objects:  new.dataset & model
# new.dataset contains the estimates of the selection effect, and selection coefficient (gamma); as well as the estimates of constraint effect (Rest) and constraint (f).
eb.res = SnIPRE(data)

res = eb.res$new.dataset
model = eb.res$model

write.table(res, file = "eb_results.csv", sep  = ",", row.names = FALSE)

#################################################################
## Part (2)  Bayesian Implementation (R2WinBUGS package,
##      	B_SnIPRE_source.R, and WinBUGS or OpenBUGS) necessary
#################################################################

source("B_SnIPRE_source.R")
source("my.jags2.R")
library(lme4)
library(R2jags)
library(arm)

test <- read.table("Snipre_Drosae_Dspinosa_Bayes_check_positive.txt", header = TRUE)  # sample data set
#BSnIPRE.run <- function(mydata, path = ".", burnin = 500, thin = 5, iter = 2500){
  # path will be where the chains are stored, and must also be where the ".bug" model is located
  # burnin, thin, and iter (number iterations after burnin) are for MCMC samples
BSnIPRE.run(test, burnin = 10000, thin = 4, iter = 15000)

# check to make sure it finished correctly:
# if a "sample" file is in your working directory (getwd()), or the path you specified)
# is empty or not there, there is a problem

load("samples")

res.mcmc <- samples

#BSnIPRE <- function(data.mcmc,mydata){
# outputs 2 objects:  new.dataset & effects
# new.dataset contains the estimates of the selection effect, and selection coefficient (gamma); as well as the estimates of constraint effect (Rest) and constraint (f).
# the "effects" may be useful if you are interested in estimation
# of population parameters (gamma, constraint) with other assumptions than the PRF

b.res <- BSnIPRE(res.mcmc, test)

bres = b.res$new.dataset

write.table(bres, file = "bayesian_results.csv", sep  = ",", row.names = FALSE)
```

**Code S3**. RNAseq analysis

*Quality control:*

fastqc file.fastq.gz

*Alignment:*

STAR --runThreadN 20 --runMode genomeGenerate --genomeDir Alignments --genomeFastaFiles /home/ksenia/assembly.fasta --sjdbGTFfile /home/ksenia/gene_prediction.gtf --sjdbOverhang 149 --genomeSAindexNbases 13

STAR --runThreadN 10 --genomeDir Alignments --readFilesCommand zcat --readFilesIn R1.fq.gz R2.fq.gz --chimOutType withinBAM --outSAMtype BAM SortedByCoordinate --outFileNamePrefix Out

*Genome annotation:*

braker.pl --genome=assembly.fasta --bam=RNAseq.out.bam --gff3 --useexisting

*Feature counts:*

featureCounts RNAseq.out.bam -a gene_prediction.gff3 -T 5 -M -p -t gene -o gene_counts

*Differential expression analysis (Love et al. 2014):*

*The pipeline is available at:* [*http://bioconductor.org/packages/devel/bioc/vignettes/DESeq2/inst/doc/DESeq2.html*](http://bioconductor.org/packages/devel/bioc/vignettes/DESeq2/inst/doc/DESeq2.html)

```
#DESeq2 R package
library(DESeq2)
library(pheatmap)
library(vsn)
library(RColorBrewer)
library(remotes)
library(ggpubr)
library(ggplot2)

#Upload the count matrix
cts <- as.matrix(read.table("C:/cygwin64/home/Ksenia/DESeq2_gene_matrix.txt", header=T))
cts0 <- cts[,c("Egg_1", "Egg_3", "Egg_4", "Larva_July_1", "Larva_July_2", "Larva_July_3", "Larva_July_4", "Larva_Sept_1", "Larva_Sept_2", "Larva_Sept_3", "Larva_Sept_4", "Gland_Oct_1", "Gland_Oct_2", "Gland_Oct_3", "Gland_Oct_4", "Gland_Nov_1", "Gland_Nov_2", "Gland_Nov_3", "Gland_Nov_4", "Head_1", "Head_2")]

#Upload sample type (factor) matrix
coldata <- read.table("C:/cygwin64/home/Ksenia/DESeq2_coldata.txt", header=T, row.names = 1)
coldata0 <- coldata[c("Egg_1", "Egg_3", "Egg_4", "Larva_July_1", "Larva_July_2", "Larva_July_3", "Larva_July_4", "Larva_Sept_1", "Larva_Sept_2", "Larva_Sept_3", "Larva_Sept_4", "Gland_Oct_1", "Gland_Oct_2", "Gland_Oct_3", "Gland_Oct_4", "Gland_Nov_1", "Gland_Nov_2", "Gland_Nov_3", "Gland_Nov_4", "Head_1", "Head_2"),]
coldata0 <- data.frame(coldata0)
coldata0$coldata0 <- factor(coldata0$coldata0)

#Sample distance matrix and visualisation
dds0 <- DESeqDataSetFromMatrix(countData = cts0, colData = coldata0, design = ~ coldata0)
ntd0 <- normTransform(dds0)
sampleDists <- dist(t(assay(ntd0)))
sampleDistMatrix <- as.matrix(sampleDists)
rownames(sampleDistMatrix) <- paste(ntd0$coldata0, sep="-")
colnames(sampleDistMatrix) <- paste(ntd0$coldata0, sep="-")
colors <- colorRampPalette(brewer.pal(9, "PuBuGn"))(255)
pheatmap(sampleDistMatrix,
         clustering_distance_rows=sampleDists,
         clustering_distance_cols=sampleDists,
         col=colors)

#Count matrices for pairwise comparisons
cts1 <- cts[,c("Larva_July_1", "Larva_July_2", "Larva_July_3", "Larva_July_4", "Egg_1", "Egg_3", "Egg_4")]

#Factor matrices for pairwise comparisons
coldata1 <- coldata[c("Egg_1", "Egg_3", "Egg_4", "Larva_July_1", "Larva_July_2", "Larva_July_3", "Larva_July_4"),]
coldata1 <- data.frame(coldata1)
coldata1$coldata1 <- factor(coldata1$coldata1)

#Create the object dds from the matrices for further differential expression analysis
dds1 <- DESeqDataSetFromMatrix(countData = cts1, colData = coldata1, design = ~ coldata1)
#log2(n + 1) transformation of the counts for the principal component analysis
ntd1 <- normTransform(dds1)

#Principal Component Analysis
pca1 <- plotPCA(ntd1,intgroup = "coldata1")
pca1

#Differential expression analysis. The function DESeq comprises (1) estimation of size factors, (2) estimation of dispersion, and negative binomial generalized linear model fitting for the log2 fold changes in gene counts (see in detail Love et al. 2014)
dds1 <- DESeq(dds1)

#Plotting dispersion and assessing model fitting
plotDispEsts(dds1)

#Show significant (adjusted p-value < 0.05) log2 fold changes (<0 for down-regulated genes and  >0 for up-regulated genes)
res1 <- results(dds1)
summary(res1, alpha=0.05)

#Visualise log2 fold changes
plotMA(res1, alpha=0.05)
```

*Bidirectional best hits:*

tblastn -query putative_larva_venom_genes_Drosae.aa -db Biorhiza_transcriptome.fa -outfmt 6

Choose query_seq_ids and subject_seq_ids with the highest bit score from tblastn output (List 1)

makeblastdb -in putative_larva_venom_genes_Drosae.aa -dbtype prot -parse_seqids

blastx -query Biorhiza_best_hit_seq.fa -db putative_larva_venom_genes_Drosae.aa -outfmt 6

Choose query_seq_ids and subject_seq_ids with the highest bit score from blastx output (List 2).

Compare List 1 and List 2 and choose the genes showing the same alignments.
