## Supplementary material for "Transcriptome analysis of *Diplolepis rosae*: revealing overexpression of genes potentially associated with insect immune response and gall formation at early larval stages": Figures

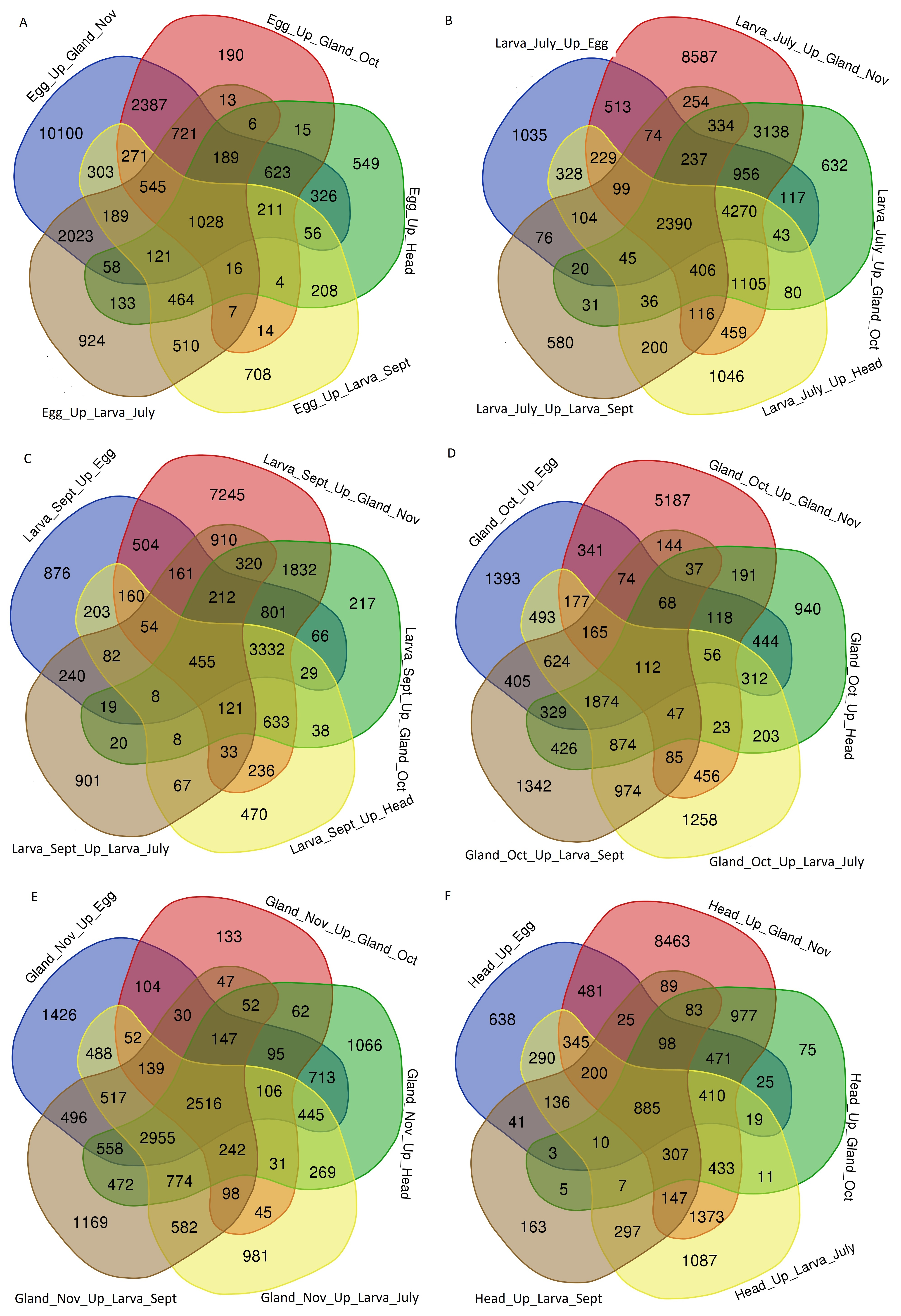

**Figure 1. Number of genes specifically expressed in the *Diplolepis rosae* (A) eggs removed from a female adult (Egg_Up), (B) mid-July larva (Larva_July_Up), (С) early September larva (Larva_Sept_Up), (D) October larva salivary gland (Gland_Oct_Up), (E) November larva salivary gland (Gland_Nov_Up), and (F) female adult head (Head_Up) found by relative differential gene expression analysis.** Up, relatively over-expressed genes found in the first sample compared to the second one. Venn diagrams were drawn by the tool available at <https://bioinformatics.psb.ugent.be/webtools/Venn/>.

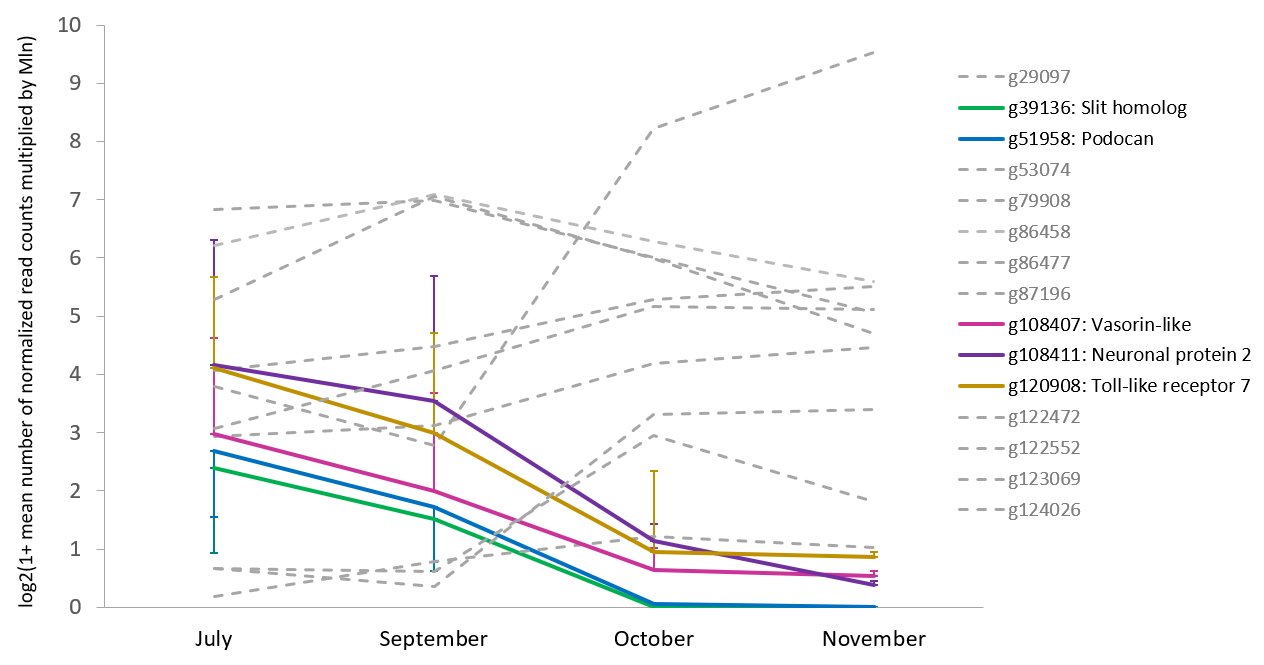

**Figure 2. Gene expression dynamics of proteins containing leucine-rich repeats in *Diplolepis rosae*.** Results are presented as mean ± SEM. Genes g108407, g108411, and g120908 show only the upper SEM bound. Genes g39136 and g51958 show only the lower SEM bound. Colored lines represent genes with significantly higher expression in July and September (combined sample ‘mid-July *D. rosae* larva + early September *D. rosae* larva + mid-July *D. eglanteriae* larva’) compared to October and November (‘October *D. rosae* larva salivary glands + November *D. rosae* larva salivary glands’). Dashed grey lines represent genes that did not show significant overexpression in July and September compared to October and November. g29097, slit homolog; g53074, chaoptin-like protein; g79908, U2 small ribonucleoprotein A’; g86458, peroxidasin-like protein; g86477, protein phosphatase 1 regulatory subunit 42-like; g87196, chaoptin; g122472, slit-like protein; g122552, insulin-like growth factor-binding protein; g123069, fibronectin type III domain containing protein; g124026, follicle-stimulating hormone receptor.

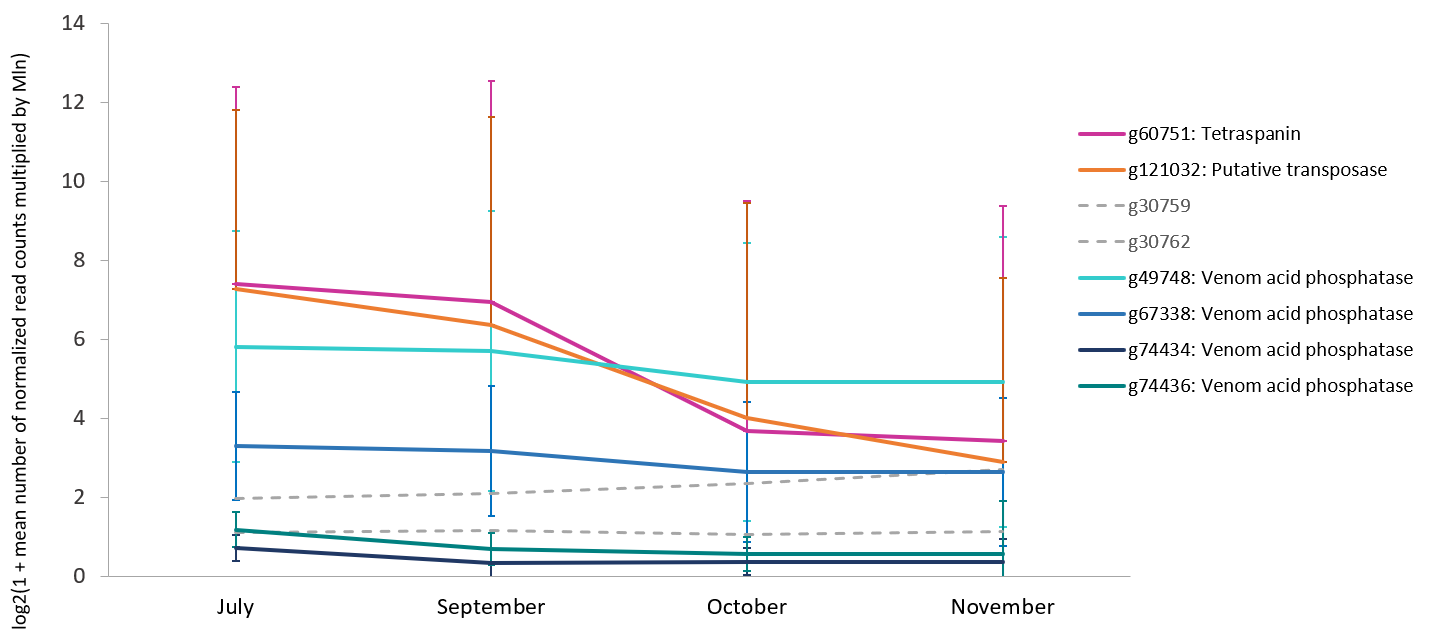

**Figure 3. Expression of genes up-regulated in the venom gland and the larvae of *Diplolepis rosae*.** Results are presented as mean ± SEM. Genes g60751 and g121032 show only the upper SEM bound. Colored lines represent genes with significantly higher expression in July and September (combined sample ‘mid-July *D. rosae* larva + early September *D. rosae* larva + mid-July *D. eglanteriae* larva’) compared to October and November (‘October *D. rosae* larva salivary glands + November *D. rosae* larva salivary glands’). Dashed grey lines represent genes that did not show significant overexpression in July and September compared to October and November. g30759 and g30762, Bi-VSP-like venom serine protease.

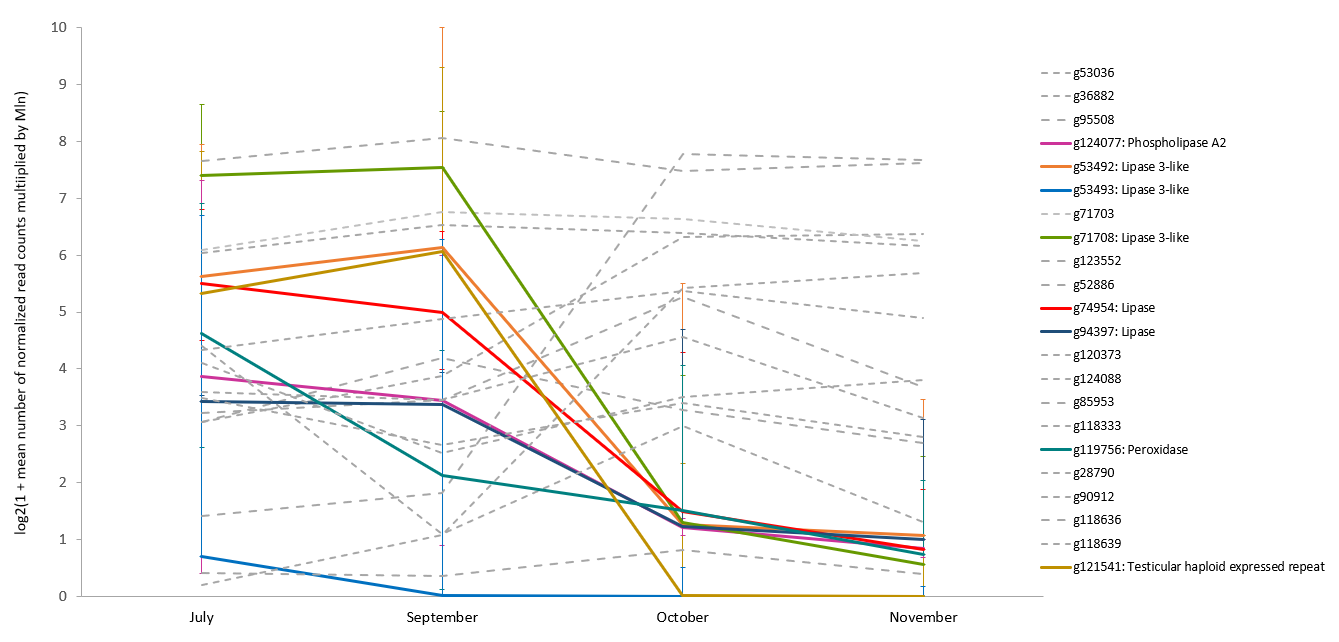

**Figure 4. Expression of *Diplolepis rosae* genes orthologous to genes up-regulated in the venom gland of *Biorhiza pallida*.** Results are presented as mean ± SEM (upper bounds). Colored lines represent genes with significantly higher expression in July and September (combined sample ‘mid-July *D. rosae* larva + early September *D. rosae* larva + mid-July *D. eglanteriae* larva’) compared to October and November (‘October *D. rosae* larva salivary glands + November *D. rosae* larva salivary glands’). Dashed grey lines represent genes that did not show significant overexpression in July and September compared to October and November. g53036, chitooligosaccharidolytic beta-N-acetylglucosaminidase; g36882, group XII secretory phospholipase A2; g95508, phospolipase A2-like; g71703, lipase-3-like enzyme; g123552: lipase family; g52886: lipase; g120373: pancreatic triacylglycerol lipase; g124088: peroxidase-like enzyme; g85953, peroxidase; g118333, peroxidasin homolog; g28790, protein D2-like; g90912, apyrase; g118636, inosine-uridine preferring nucleoside hydrolase; g118639, lysozyme C1-like.

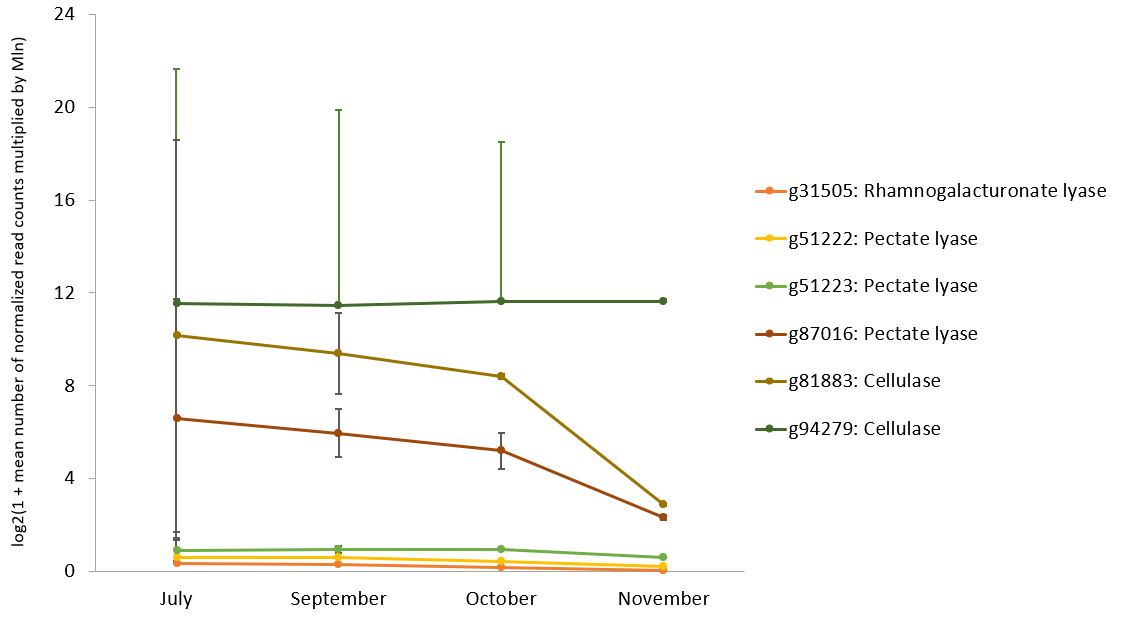

**Figure 5. Gene expression dynamics of plant cell wall degrading enzymes in *Diplolepis rosae*.** Results are presented as mean ± SEM. Gene g94279 shows only the upper SEM bound. When comparing gene expression between summer (combined sample ‘mid-July *D. rosae* larva + early September *D. rosae* larva + mid-July *D. eglanteriae* larva’) and autumn (‘October *D. rosae* larva salivary glands + November *D. rosae* larva salivary glands’), no significant results were found.

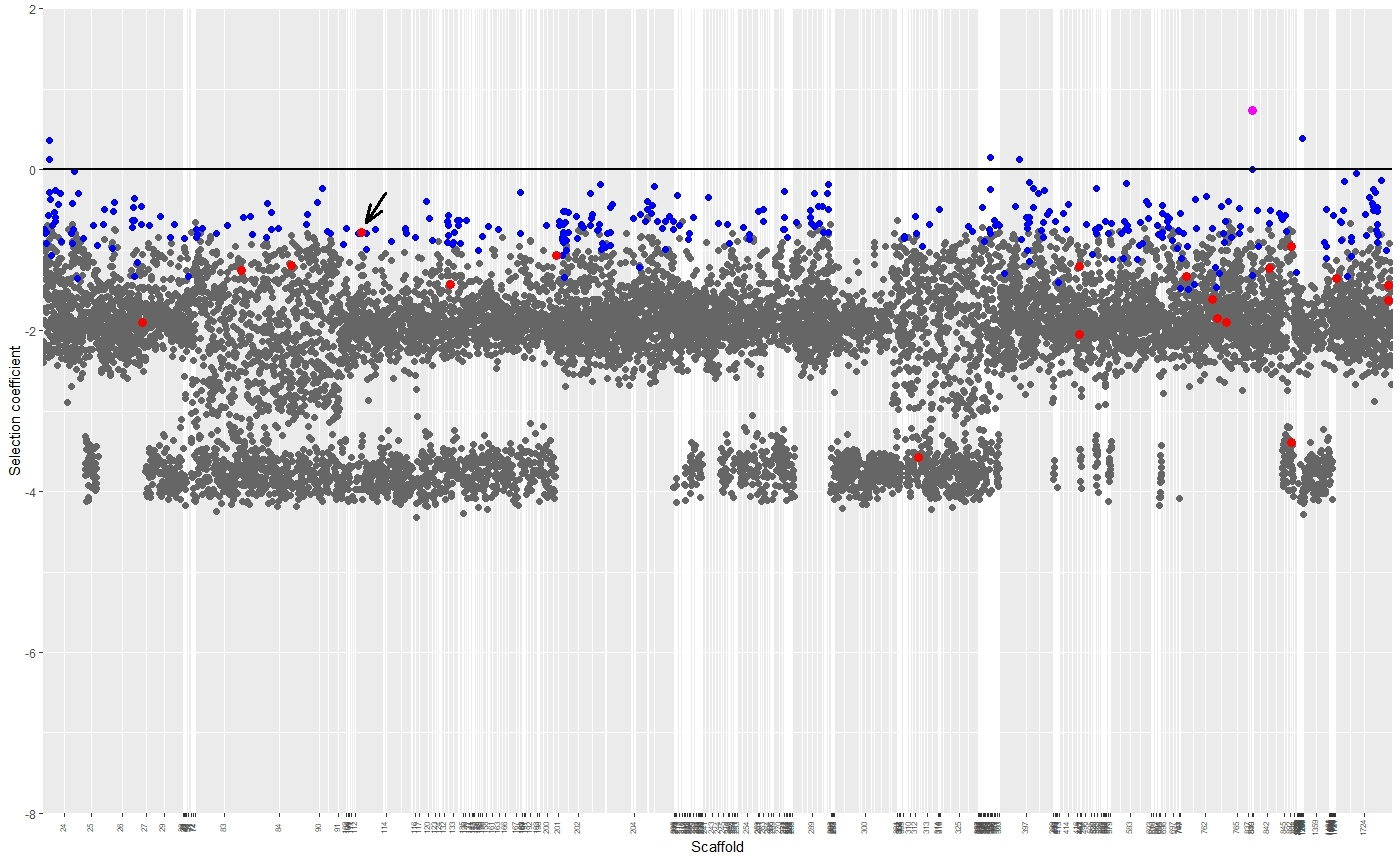

**Figure 6. Genomic scan of the selection coefficient (s) estimated by SnIPRE (Eilertson et al. 2012) in the protein-coding sequences of the *Diplolepis rosae* genome.** The numbers indicate scaffolds in the genome assembly. The black line indicates the selection coefficient equal to zero, which reflects the absence of selection. Dark grey points: mean s estimations corresponding to genes under negative selection. Blue points: mean s estimations corresponding to genes under neutral selection. Rose point: mean s estimation corresponding to a gene under positive selection. Red points: genes that were found to be up-regulated in summer *D. rosae* larvae compared to autumn *D. rosae* larvae (combined sample ‘mid-July *D. rosae* larva + early September *D. rosae* larva + mid-July *D. rosae* larva’ vs combined sample ‘October *D. rosae* salivary gland + November *D. rosae* salivary gland’) (**Fig. 2-4**). All these genes are under negative selection except for that encoding tetraspanin (g60751, marked by flash), that is under neutral selection.
